# Stathmin 1 regulates mitophagy and cellular function in hematopoietic stem cells

**DOI:** 10.1101/2025.03.10.642434

**Authors:** Luana Chiquetto, Meg Schuetz, Qian Dong, Mia Warmka, Liana Valin, Ashley Jones, Patrick Hunt, Copper Petermeier, Jie Wang, Nate Roundy, Zev J. Greenberg, Wei Yang, Christine R. Zhang, Grant A. Challen, Cliff J. Luke, Robert A. J. Signer, Wandy L. Beatty, Stephen Sykes, Weikai Li, David J. Kast, Laura G. Schuettpelz

**Affiliations:** Department of Pediatrics, Washington University School of Medicine, St. Louis, MO 63110, USA; Department of Cell Biology and Physiology, Washington University School of Medicine, St. Louis, MO 63110, USA; Department of Genetics, Washington University School of Medicine, St. Louis, MO 63110, USA; Department of Medicine, Washington University School of Medicine, St. Louis, MO 63110, USA; Department of Medicine, Sanford Stem Cell Institute, Moores Cancer Center, University of California at San Diego, La Jolla, CA 92093, USA; Department of Molecular Microbiology, Washington University School of Medicine, St. Louis, MO 63110, USA; Department of Biochemistry and Molecular Biophysics, Washington University School of Medicine, St. Louis, MO 63110, USA

**Keywords:** stathmin 1, hematopoietic stem cells, metabolism, autophagy, microtubules

## Abstract

Stathmin 1 is a cytoplasmic phosphoprotein that regulates microtubule dynamics via promotion of microtubule catastrophe and sequestration of free tubulin heterodimers. Stathmin 1 is highly expressed in hematopoietic stem cells (HSCs), and overexpressed in leukemic cells, however its role in HSCs is not known. Herein, we found that loss of Stathmin 1 is associated with altered microtubule architecture in HSCs, and markedly impaired HSC function. Transcriptomic studies suggested alterations in oxidative phosphorylation in *Stmn1*^*-/-*^ HSCs, and further mechanistic studies revealed defective mitochondrial structure and function in the absence of Stathmin 1 with increased ROS production. Microtubules associate with mitochondria and lysosomes to facilitate autophagosome formation and mitophagy, and indeed we found that this critical mitochondrial quality control process is impaired in Stathmin 1-deficient HSCs. Finally, stimulation of autophagy improved the colony forming ability of *Stmn1*^*-/-*^ hematopoietic stem and progenitor cells. Together, our data identify Stathmin 1 as a novel regulator of mitophagy and mitochondrial health in HSCs.

**Key Points:**

- The microtubule regulating protein Stathmin 1 is highly expressed in HSPCs and promotes normal microtubule architecture.
- Loss of Stathmin 1 in HSPCs leads to impaired autophagy with abnormal mitochondrial morphology, decreased respiratory capacity, and impaired cellular function.

## Introduction

Stathmin 1 is a cytoplasmic phosphoprotein that regulates microtubule dynamics via directly promoting microtubule catastrophe or by regulating microtubule assembly through the sequestration of alpha/beta tubulin heterodimers.^1-3^ Also known as Oncoprotein 18 (OP18) or Leukemia-associated phosphoprotein p18 (LAP18), Stathmin 1 is highly expressed in a wide variety of tumors, including myeloid and lymphoid leukemias, and its down-regulation can promote apoptosis and diminish cell proliferation and migration.^4-8^

In the normal hematopoietic system, Stathmin 1 is highly expressed in hematopoietic stem cells (HSCs) and progenitors, and its expression declines with cellular differentiation.^9^ Several studies have implicated Stathmin 1 in the regulation of hematopoietic lineage maturation.^10^ For example, Stathmin 1 expression is important for normal erythroid differentiation,^11^ and inhibition of Stathmin 1 in erythroleukemic cells inhibits their differentiation in response to hemin. In addition, downregulation of Stathmin 1 expression is important for normal megakaryocyte maturation and platelet production.^12^ A recent study by Ramlogan-Steele and colleagues^13^ showed that *Stmn1*^*-/-*^ mice have reduced erythrocyte progenitors and a mild, progressive macrocytic anemia that persists throughout life. Megakaryocytes (MK) and platelets are also altered in the *Stmn1*^*-/-*^ mice. Early on (at age 3 weeks), these mice have reduced megakaryocytes in the marrow and spleen, with reduced circulating platelet counts. As they age, however, platelet counts increase above normal controls, and the bone marrow MKs of aged *Stmn1*^*-/-*^ mice display enlarged size and ploidy compared to their normal counterparts.^13^

While prior studies have established a role for Stathmin 1 in normal erythro-megakaryopoiesis, its role in hematopoiesis remains unclear. Specifically, the role of Stathmin 1 in regulating HSC function is not known. Given the high expression of Stathmin 1 in HSCs, as well as the importance of microtubule dynamics to hematopoietic stem and progenitor cell differentiation,^14^ we hypothesized that loss of Stathmin 1 may impact normal HSC function.

Indeed, we found that while *Stmn1*^*-/-*^ mice have normal numbers of immunophenotypic HSCs, their function is markedly impaired in competitive transplants. Further, *Stmn1*^*-/-*^ mice have delayed hematopoietic recovery from the chemotherapeutic agent 5-fluorouracil (5-FU). Consistent with its role as a regulator of microtubule dynamics, *Stmn1*^*-/-*^ HSCs have altered microtubule networks. Transcriptome analysis of *Stmn1*^*-/-*^ HSCs suggested mitochondrial metabolism defects, and subsequent metabolic studies confirmed that *Stmn1*^*-/-*^ HSCs have impaired mitochondrial respiratory capacity and increased reactive oxygen species (ROS) production. Further, *Stmn1*^*-/-*^ HSCs have abnormal mitochondrial morphology, as revealed by transmission electron microscopy (TEM). Microtubule-mitochondria associations facilitate mitophagy, an essential process for regulating of mitochondria homeostasis, and indeed this process is impaired in the absence of Stathmin 1. Together, these data suggest a model whereby Stathmin 1 regulates microtubule dynamics in HSCs to support mitochondrial quality control and cellular function.

## Methods

### Mice

*Stmn1*^*-/-*^ (B6.129P2-*Stmn1*^*tm1Wed*^/J),^15^ C57BL/6J (CD45.2), and C57BL/6 mice (B6.SJL-Ptprc^a^ Pepc^b^ /BoyJ) carrying the CD45.1 allele were purchased from the Jackson Laboratory (Bar Harbor, ME). WT littermates were used throughout the study as controls. Young adult (8-10 week) sex- and age-matched mice were used for all experiments in accordance with the guidelines of the Washington University Animal Studies Committee.

### Flow cytometry

Peripheral blood was obtained by retro-orbital sampling, and cell counts determined using an Element HT5 (Heska Corporation, Loveland, CO, USA) automated cell counter. Bone marrow cells were isolated by centrifugation of long bones at 6,000 rpm for 3 minutes. After lysis of red blood cells, samples were stained (see Supplemental Table 1 for list of antibodies) and analyzed on NovoCyte flow cytometer (Agilent Technologies, San Diego, CA, USA) or an Aurora flow cytometer (Cytek Biosciences, Freemont, CA, USA). Data was analyzed with FlowJo software (version 10.5.3; TreeStar, Ashland, OR, USA).

### Long term repopulating assays

Whole bone marrow (WBM) cells from *Stmn1*^*-/-*^ mice or WT littermates (CD45.2) were transplanted with or without WT (CD45.1) competitor WBM cells (2□×□10^6^ cells total) via retro-orbital injection into lethally irradiated (two doses of 550 cGy spaced 4h apart) WT (CD45.1/CD45.2) recipients. For intra-tibial transplants, a total of 2.5□×□10^5^ cells were transplanted directly into the tibias of lethally irradiated recipients. Peripheral blood chimerism was determined via retro-orbital sampling every 6 weeks.

### Seahorse Mito Stress and Glycolysis Stress Tests

Seahorse Mito Stress Test and Glycolysis Stress Test kits (Agilent Technologies, Santa Clara, CA, USA) were used to measure oxygen consumption rate (OCR) and extracellular acidification rate (ECAR) using a 96-well Seahorse Bioanalyzer XFe96 following manufacturer’s instructions. c-Kit+ cells were plated at a density of 1×10^5^ cells per well into 96 well plates with StemSpan media with thrombopoietin (TPO; 100 ng/mL, PeproTech, Cranbury, NJ, USA), stem cell factor (SCF; 100 ng/mL, PeproTech), IL-3 (10 ng/mL, PeproTech), FLT3L (50 ng/mL, PeproTech), and G-CSF (10ng/mL, PeproTech) for 18 hours at 37C with 5%CO_2_. For Mito Stress Test (OCR), cells were then washed and resuspended in XF base media (RPMI supplemented with Glutamine (200 mM), glucose (2.5 M), and Sodium-pyruvate (100 mM)). Plates were then spun for 5 minutes at 1600 rpm. The injection port A on the sensor cartridge was loaded with 1μM Oligomycin (Oligo), port B was loaded with 2μM FCCP and port C was loaded with 0.5μM Rotenone/Antimycin (ROT/AA). For sensor calibration, cells were incubated at 37C in a non-CO2 incubator, then plate was immediately placed onto calibrated XF96 extracellular flux analyzer for the Mito Stress test. For the Glycolysis Stress Test (ECAR) cells were resuspended in XF base media supplemented with L-glutamine (200nM). The injection port A on the sensor cartridge was loaded with 10mM glucose, port B with 2μM oligomycin and 50mM 2-DG was loaded in port C. Plate was incubated in 37C non-CO2 incubator during sensor calibration. Then, plate was loaded into calibrated XF96 extracellular flux analyzer for glycolysis stress test.

### Immunofluorescence staining, imaging and analysis

HSCs were sorted in Opti-MEM media onto poly-L-lysine coated slides, then incubated for one hour at 4C. Cells were then fixed with 4% paraformaldehyde for 20 minutes at room temperature (RT), permeabilized with 0.1% triton X-100 and blocked with 2% bovine serum albumin. Cells were immuno-stained with Drp1(DNM1L) monoclonal antibody (1:100, Invitrogen #MA26255), Parkin polyclonal antibody (1:100, Invitrogen #PA13399), LC3B polyclonal antibody (1:100, Invitrogen #PA5-32254), Tom20 mouse monoclonal antibody (1:100, Santa Cruz, #sc17764), or tubulin-alpha monoclonal antibody (1:100, BioLegend, #627910) in blocking buffer at RT for one hour. Secondary staining in AF488 goat anti-mouse (Invitrogen, #A11001) and AF568 donkey anti-rabbit (Invitrogen, #A10042) were done after washes and incubated at RT for one hour. Slides were mounted using anti-fade gold reagent with DAPI (Vectashield, #H-1200-10). Images were acquired with a Zeiss LSM 880 confocal microscope equipped with Airyscan Super Resolution Imagine module using a 63X/1.46 Alpha Plan Apochromat objective lens (Zeiss MicroImaging, Jena, Germany). Integrated fluorescence intensity density of images was calculated using Fiji software on unadjusted raw images. Background subtraction and adjustment of brightness were done on all sets of images avoiding low-end and high-end clipping of data. Colocalization measurements were done on Volocity software using the colocalization feature. Puncta counts were done on Volocity software using the count objects feature, separating objects bigger than 3μm.

To determine mitophagy flux, HSCs were sorted into StemSpan SFEM media with TPO (100 ng/mL) and SCF (100 ng/mL), and Leupeptin (100 μM, ThermoFisher, Waltham, MA, USA) was added to the media for two hours. Cells were then plated onto poly-L-lysine slides, allowed to rest for 1 hour prior to fixation, permeabilization and LC3 immunostaining as described above. Mitophagy flux was calculated subtracting the baseline levels of LC3 and leupeptin treated LC3 integrated density levels.

### Statistical analysis

Data are presented as mean ± standard error of the mean (SEM), unless otherwise stated. In some cases, the data were log-transformed when the data skewed prior to statistical analysis. Statistical significance was assessed using an unpaired two-tailed Student’s t-test or two-way analysis of variance (ANOVA) for untransformed data and an unpaired Student’s t-test with Welch’s correction for log-transformed data. GraphPad Prism (Version 9.3.1) was used for all statistical analyses (GraphPad Software, La Jolla, CA, USA). In all cases, *p<0.05, **p<0.01, ***p<0.001, and ****p<0.0001.

### Data sharing

RNA-seq data are available at GEO under accession number XXXX (To be deposited upon acceptance)

## Results

### Stmn1^-/-^ HSPCs have an altered microtubule network

To assess the role of Stathmin 1 in HSCs, we first visualized Stathmin 1 protein expression and localization by confocal microscopy. Consistent with previous reports of high RNA expression in HSCs,^9^ we detected Stathmin 1 protein in sorted HSCs (Lineage-c-Kit+ Sca-1+ CD150+ CD48-cells) from young adult (8-10 week) wild-type (WT) mice (Supplemental Figure 1A). Further, in accordance with its known role as a microtubule destabilizer,^2^ Stathmin 1 expression in HSCs largely co-localizes with tubulin (Supplemental Figure 1B). To assess the effects of Stathmin 1 loss, we next examined the microtubule network in sorted c-Kit+ hematopoietic stem and progenitor cells (HSPCs) from *Stmn1*^*-/-*^ mice and WT controls. For these experiments, we used a methanol fixation approach to better visualize the microtubule network. Confocal imaging of tubulin revealed several differences between the microtubules in *Stmn1*^*-/-*^ HSPCs compared to WT. Consistent with prior reports, microtubules are distributed in a polar fashion in the WT HSPCs, with fine microtubule networks emanating from the microtubule organizing center (Figure 1A, left panels).^14,16^ In contrast, *Stmn1*^*-/-*^ HSPCs display thickened microtubules (Figure 1A, right panels). In addition, tubulin levels as determined by mean grey value are reduced in *Stmn1*^*-/-*^ HSPCs, and *Stmn1*^*-/-*^ HSPCs are significantly smaller than WT (Figure 1B-D). Thus, Stathmin 1 is expressed in HSCs, and the microtubule architecture is altered in its absence.

**Figure 1.**
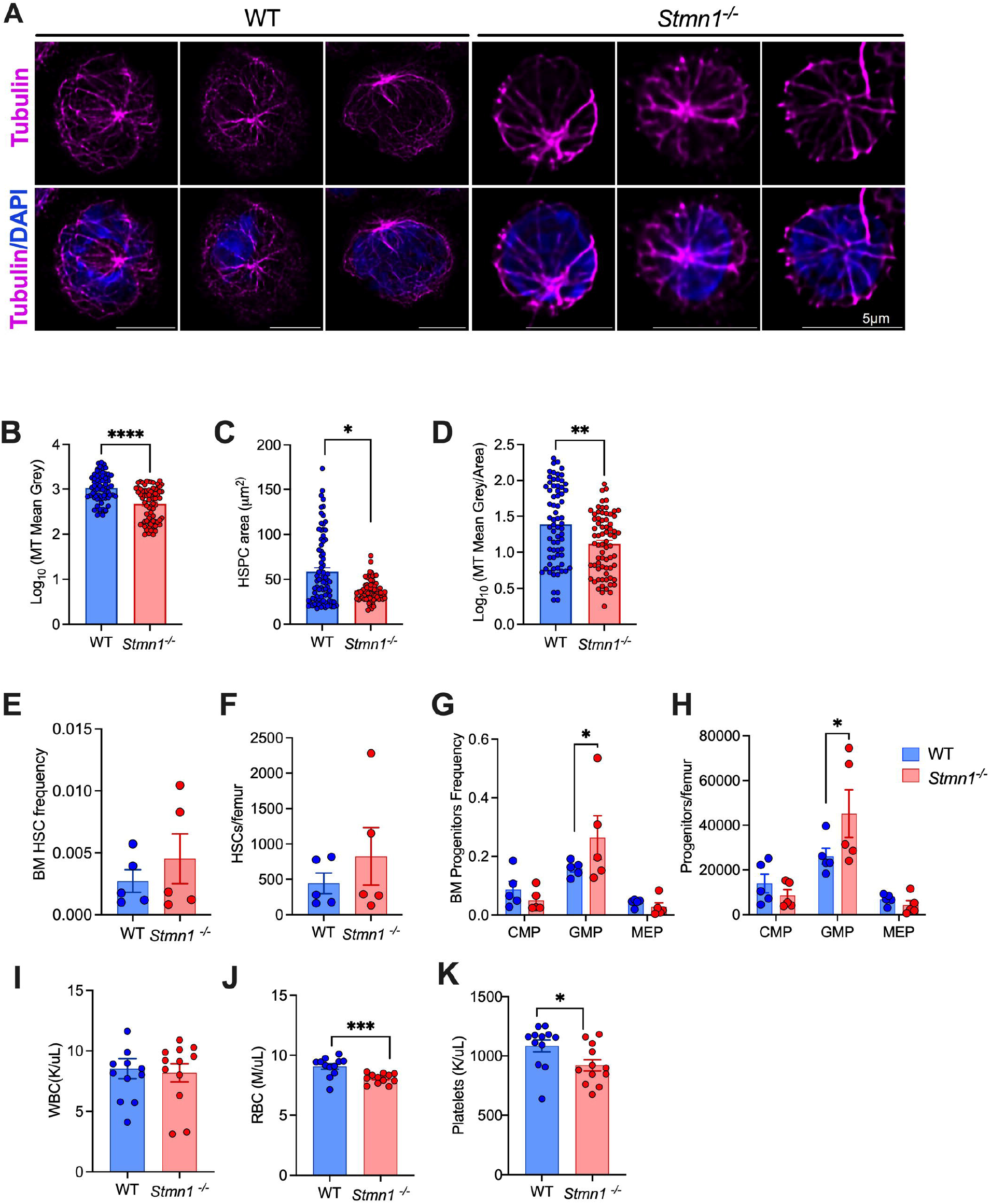
*Stmn1*^*-/-*^ HSPCs have an altered microtubule network. (A) Representative images of the microtubule network (see Supplemental Methods) in HSPCs (c-Kit+ cells) from WT and *Stmn1*^*-/-*^ mice. Microtubules were stained with alpha-beta tubulin (magenta) and DAPI (blue). (B-D) Quantification of tubulin intensity (B), HSPC area (C) and the ratio of tubulin intensity to area (D). (E-H) Shown are bone marrow HSC frequency (E) and absolute numbers (F) from young adult (8–10-week-old) WT and *Stmn1*^*-/-*^ mice, as well as myeloid progenitor frequencies (G) and absolute numbers (H). See Supplemental Figure 2A for HSC and progenitor gating strategy. (I-K) Peripheral blood counts from young adult *Stmn1*^*-/-*^ mice show equivalent numbers of WBCs (I), and modest decreases in RBCs (J) and platelets (K) compared to WT. n=5-12 mice per group over 3 independent experiments. Error bars represent mean +/-SEM. *p<0.05, ***p<0.001 by unpaired Student’s t-test (E-L), Mann-Whitney (C) or Welch’s t-test (B, D).

### Stmn1^-/-^ HSCs have impaired function

To further address the specific role of Stathmin 1 in HSCs, we assessed HSC numbers and function in *Stmn1*^*-/-*^ mice. Young adult (8-10 week) *Stmn1*^*-/-*^ mice have no significant differences in the frequency or absolute numbers of HSCs in the bone marrow compared to WT controls (Figure 1E-F). However, there is a modest increase in granulocyte-monocyte progenitors (Figure 1G-H), indicating a potential role of Stathmin 1 in the differentiation of these progenitors. Peripheral blood analysis showed, in accordance with prior reports, that *Stmn1*^*-/-*^ mice have mild anemia and thrombocytopenia compared to WT littermates (Figure 1I-K).^12,13^

Next, we assessed HSC function using competitive transplants. Whole bone marrow (WBM) cells from *Stmn1*^*-/-*^ mice or their WT littermates (CD45.2) were mixed 1:1 with WBM from WT competitor (CD45.1), transplanted into lethally irradiated WT recipients (CD45.1/45.2), and peripheral blood chimerism was monitored over time (Figure 2A). Despite having normal HSC numbers (Figure 1E-F), the loss of Stathmin 1 led to a marked reduction in multilineage repopulating activity, with approximately 80% decrease in peripheral blood chimerism of *Stmn1*^*-/-*^ cells compared to WT (Figure 2B-C). Bone marrow chimerism of *Stmn1*^*-/-*^ cells (both total WBCs and HSCs) was similarly reduced at 24 weeks post-transplant (Figure 2D-E). To test self-renewal, bone marrow from primary recipients was transplanted into secondary recipients after 24 weeks. As shown in Supplemental Figure 3A-B, *Stmn1*^*-/-*^ bone marrow did not engraft upon secondary transplantation. Importantly, bone marrow homing was not significantly different between WT and *Stmn1*^*-/-*^ c-Kit+ HSPCs (Figure 2F), and similar engraftment results were obtained when donor cells were transplanted intra-tibially (Supplemental Figure 3C-G), suggesting that the functional defect was not due to impaired homing.

**Figure 2.**
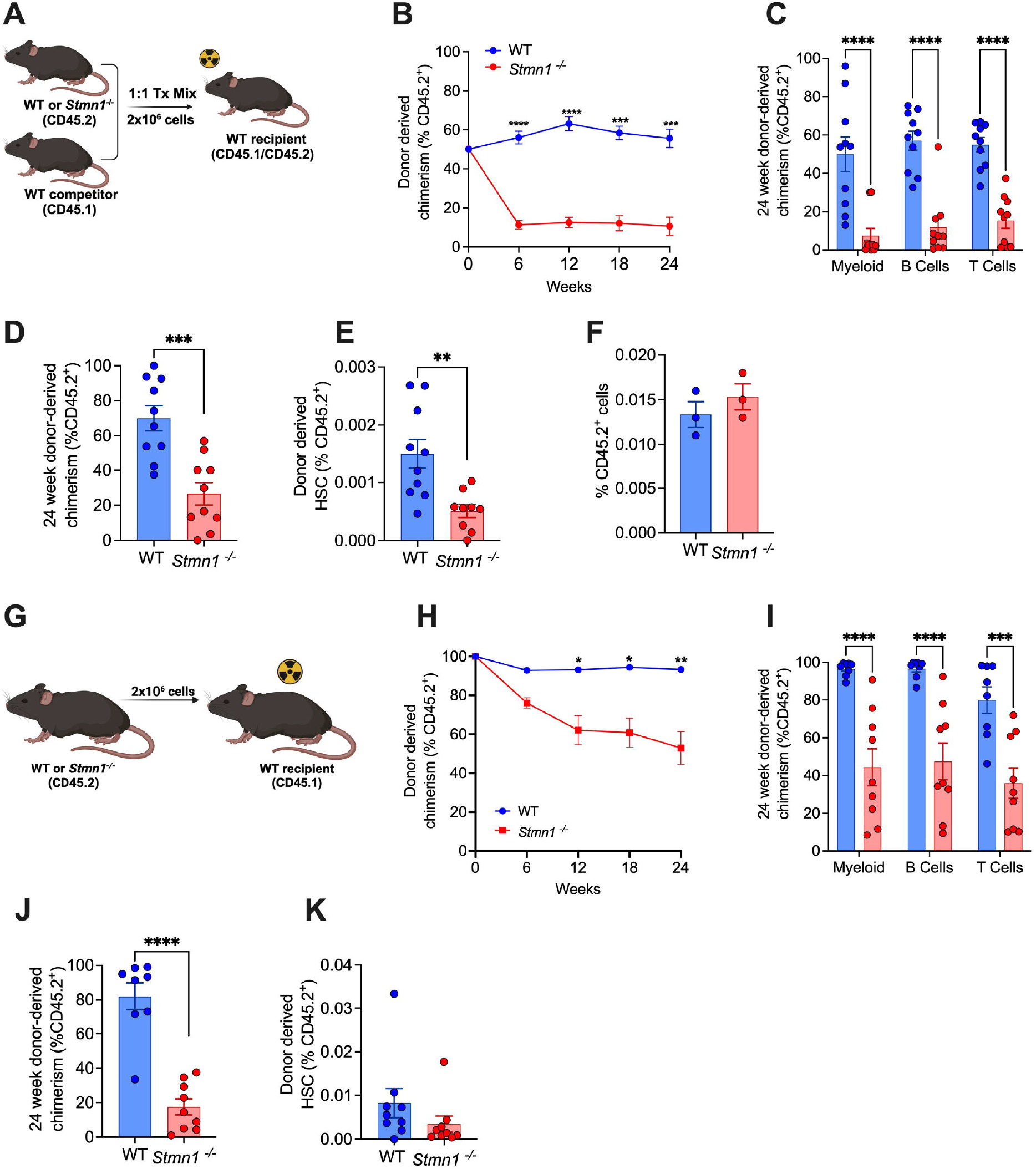
*Stmn1*^*-/-*^ HSCs have impaired repopulating activity in transplantation assays. (A) Outline of whole bone marrow (WBM) competitive transplants. WT or *Stmn1*^*-/-*^ WBM cells (CD45.2) were transplanted in a 1:1 ratio with WT competitor (CD45.1) WBM into irradiated WT recipients (CD45.1/CD45.2). Chimerism was assessed by peripheral blood (PB) flow cytometry analysis every 6 weeks (B). PB engraftment of individual lineages at 24 weeks post-transplant is shown in (C), and total bone marrow chimerism (D), and HSC chimerism at 24 weeks post-transplant is shown in (E). n = 10 recipients per group over 2 independent transplants. (F) Homing was assessed by transferring LSK cells (Lin-Sca-1+ c-Kit+ cells) from WT or *Stmn1*^*-/-*^ (CD45.2) mice into lethally irradiated CD45.1 WT mice and enumerating CD45.2 cells in the bone marrow 18 hours later. (G) Outline of non-competitive WBM transplants. WBM cells from WT or *Stmn1*^*-/-*^ mice (CD45.2) were transplanted without competitor cells into irradiated WT (CD45.1) recipients. Chimerism was assessed by PB flow cytometry analysis every 6 weeks (H). PB engraftment of individual lineages at 24 weeks post-transplant is shown in (I), and total bone marrow chimerism (J) and HSC chimerism at 24 weeks post-transplant is shown in (K). n = 8-9 recipients per group over 2 independent transplants. Error bars represent mean +/-SEM. *p<0.05, **p<0.01, ***p<0.001, and ****p<0.0001 by unpaired Student’s t-test or 2-way ANOVA.

**Figure 3.**
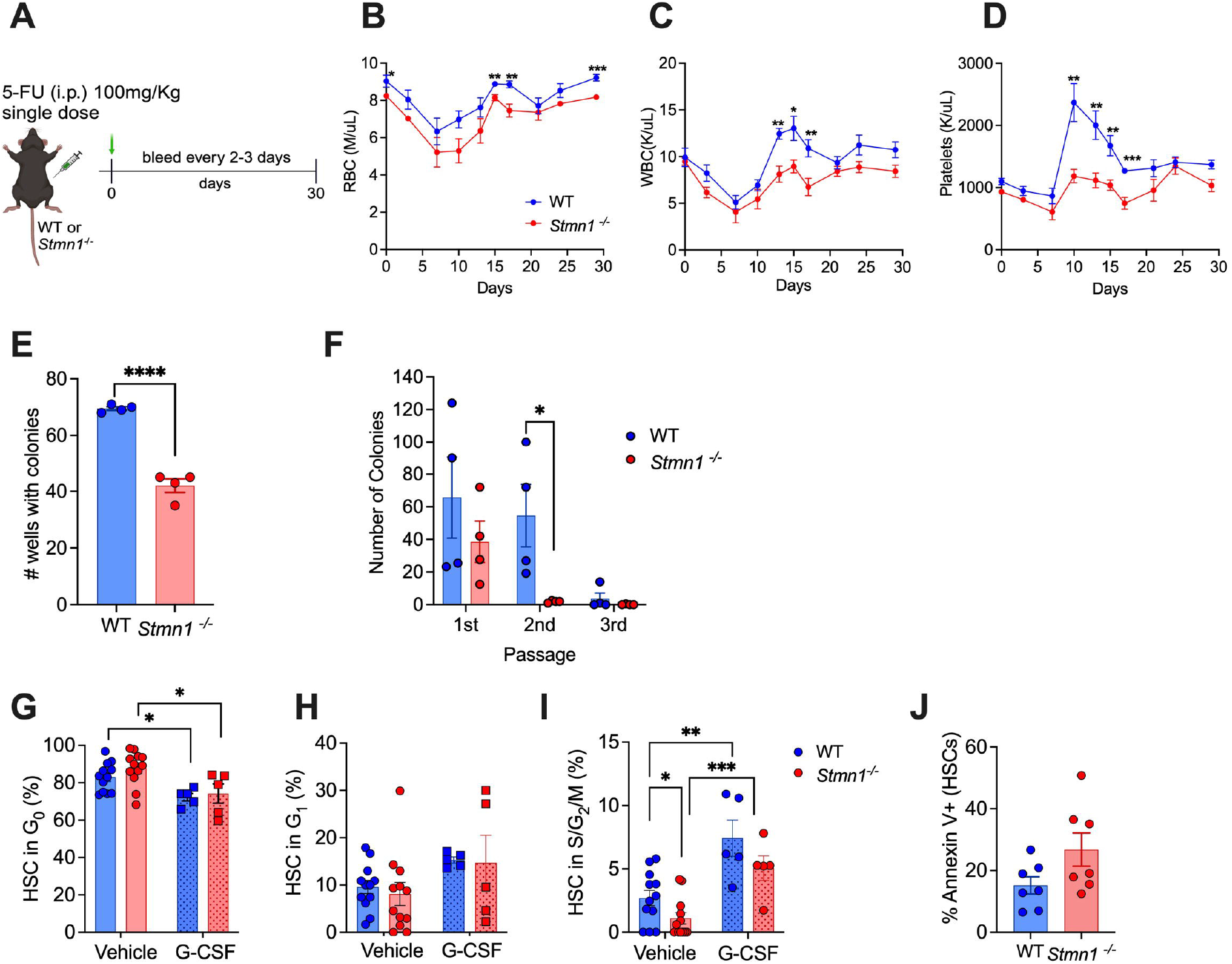
*Stmn1*^*-/-*^ HSCs have impaired function and are hyper-quiescent at baseline. (A) WT or *Stmn1*^*-/-*^ mice were treated with one single dose of 5-fluorouracil (5-FU, 100mg/kg). Peripheral blood was collected to analyze RBCs (B), WBCs (C), and platelets (D) every three days for 30 days. n=7 mice per group from two independent experiments. (E) HSCs were sorted individually into MethoCult 3434 in 96 well plates, and colonies were scored after 7 days. Shown are the total numbers of wells with colonies. (F) *Stmn1*^*-/-*^ and WT WBM cells were plated in MethoCult 3434 (1×10^4^ cells per well), and colonies were scored after 7 days. Cells were then replated (5×10^3^ cells/well) and scored every 7 days until no colonies were formed. Data represent 4 replicates over 4 independent experiments. (G-I) Cell cycling was measured by flow cytometry using Ki-67 and DAPI in WT and *Stmn1*^*−/−*^ HSCs at baseline (vehicle treatment) and following two doses of G-CSF over 24 hours. Shown are the percentages of HSCs in G_0_ (G), G_1_ (H) and S/G_2_/M (I). n= 5-12 mice per group. See Supplemental Figure 2B for representative flow plots. (J) Shown are the percent of WT and *Stmn1*^*-/-*^ HSCs that are Annexin V+ at baseline by flow cytometry. N= 7 mice per group. Error bars represent mean +/-SEM. *p<0.05. **<p0.01. ***p<0.001. ****p < 0.0001 by an unpaired Student’s t-test or 2-way ANOVA.

In addition to competitive transplants, we performed non-competitive transplants in which 2×10^6^ WBM cells from WT or *Stmn1*^*-/-*^ mice were transplanted into lethally irradiated WT recipients and peripheral blood chimerism was followed over time (Figure 2G). As with the competitive transplants, *Stmn1*^*-/-*^ cells showed markedly decreased trilineage engraftment compared to WT (Figure 2H-I). *Stmn1*^*-/-*^ cell bone marrow chimerism (total WBCs and HSCs) was also reduced compared to WT at 24 weeks post-transplant (Figure 2J-K).

To further assess HSC function in a non-transplant context, we treated *Stmn1*^*-/-*^ and WT mice with a single dose of the chemotherapeutic drug 5-fluorouracil (5-FU) or vehicle control and measured peripheral blood recovery every 3 days over 30 days. *Stmn1*^*-/-*^ mice had delayed recovery of leukocytes, erythrocytes, and platelets compared to WT (Figure 3A-D), consistent with a prior report.^13^ In addition, we sorted individual HSCs from *Stmn1*^*-/-*^ and WT mice into MethoCult and found that fewer *Stmn1*^*-/-*^ HSCs formed colonies after 7 days of culture (Figure 3E). We also plated WBM cells and found that *Stmn1*^*-/-*^ cells formed fewer total colonies than WT and were not capable of serial replating (Figure 3F). Together, these data show that Stathmin 1 is necessary for normal HSC function.

### Stmn1^-/-^ HSCs are hyper-quiescent

Given the importance of microtubules to cell cycling, and the significance of cycling to HSC function,^17^ we next assessed the cell cycle status of *Stmn1*^*-/-*^ vs WT HSCs. At baseline, *Stmn1*^*-/-*^ HSCs are more quiescent than WT HSCs, with fewer cells in S/G_2_/M phase (Figure 3G-I). Although they are hyper-quiescent at baseline, *Stmn1*^*-/-*^ HSCs can be stimulated to cycle in response to stimulation with the cytokine granulocyte colony-stimulating factor (G-CSF, Fig. 3G-I). In addition, *Stmn1*^*-/-*^ HSCs also have slightly increased, albeit not significant, apoptosis at baseline compared to WT as indicated by a trend toward increased Annexin V staining (Fig. 3J), suggesting that alterations in cell cycling and survival may contribute to loss of HSC function in the absence of Stathmin 1.

### Stmn1^-/-^ HSCs have decreased mitochondrial function

To identify mechanisms underlying the loss of HSC function in the absence of Stathmin 1, we performed bulk RNA sequencing on sorted HSCs (Lineage-c-Kit+ Sca-1+ CD150+ CD48-cells) from *Stmn1*^*-/-*^ and WT mice. We identified 212 differentially expressed genes between the groups (Supplemental Table 2). Gene set enrichment analysis (GSEA) of the transcriptome data suggested alterations in metabolism in the *Stmn1*^*-/-*^ HSCs, particularly indicating altered mitochondrial metabolism with enrichment for Myc targets, mTORC1 signaling and elevated oxidative phosphorylation (Figure 4A-B). While Myc and mTOR are important for a myriad of different signaling pathways, they also play a role in regulating genes important for mitochondrial metabolism and enhanced oxidative phosphorylation in HSCs.^18-20^ Genes associated with oxidative phosphorylation are highlighted in the volcano plot in Figure 4C.

**Figure 4.**
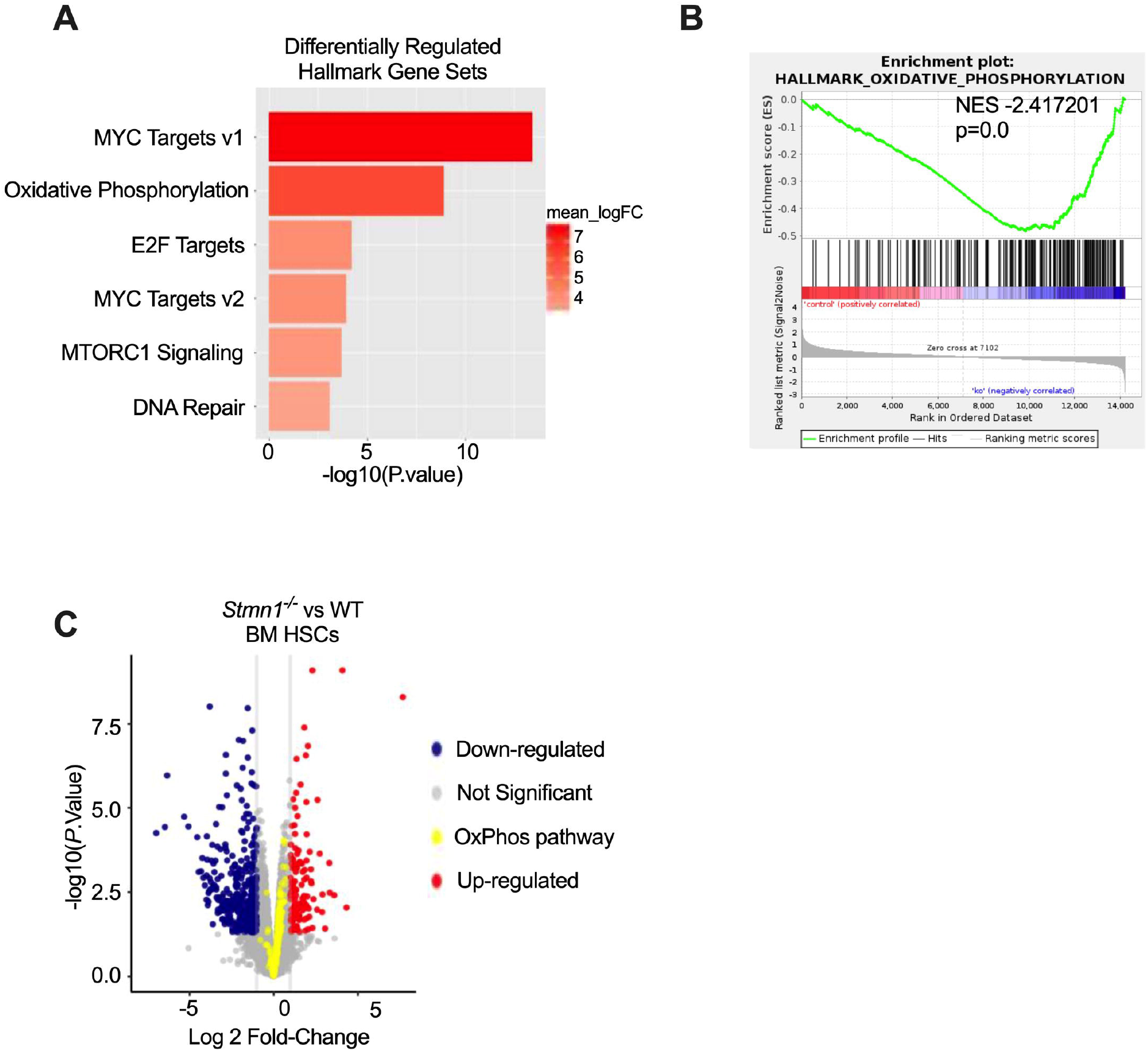
RNA-sequencing identifies altered expression of genes associated with mitochondrial metabolism in *Stmn1*^*-/-*^ HSCs. Bulk RNA sequencing was performed on sorted HSCs from *Stmn1*^*-/-*^ and WT mice. (A) Shown are the differentially expressed Hallmark Gene Sets comparing *Stmn1*^*-/-*^ to WT HSCs. (B) Enrichment plot showing significant differences in oxidative phosphorylation related genes. (C) Volcano plot of differentially expressed genes comparing *Stmn1*^*-/-*^ to WT HSCs. Oxidative phosphorylation pathway genes are highlighted in yellow.

To validate the transcriptome data suggesting alterations in mitochondrial metabolism, we next assessed mitochondrial function in the *Stmn1*^*-/-*^ and WT HSPCs (c-Kit+ cells) using the Seahorse analyzer. We found that while glycolytic capacity is similar between *Stmn1*^*-/-*^ and WT HSPCs (Supplemental Figure 4A), mitochondrial respiratory capacity is significantly reduced in the absence of Stathmin 1, with *Stmn1*^*-/-*^ cells showing decreased basal and maximal mitochondrial respiration and lower mitochondrial spare respiratory capacity (Figure 5A-B). We next assessed mitochondrial reactive oxygen species (ROS) production and mitochondrial membrane potential using the flow-based MitoSOX and TMRM (Tetramethylrhodamine methyl ester) assays and found increased ROS in *Stmn1*^*-/-*^ HSCs compared to WT (Figure 5 C-D and Supplemental Figure 4B-C). Notably, elevated ROS is associated with damaged or impaired mitochondria that lack effective mitochondrial quality control mechanisms and/or ROS control measures,^21^ and elevated ROS can be detrimental to HSC function.^19,22^

**Figure 5.**
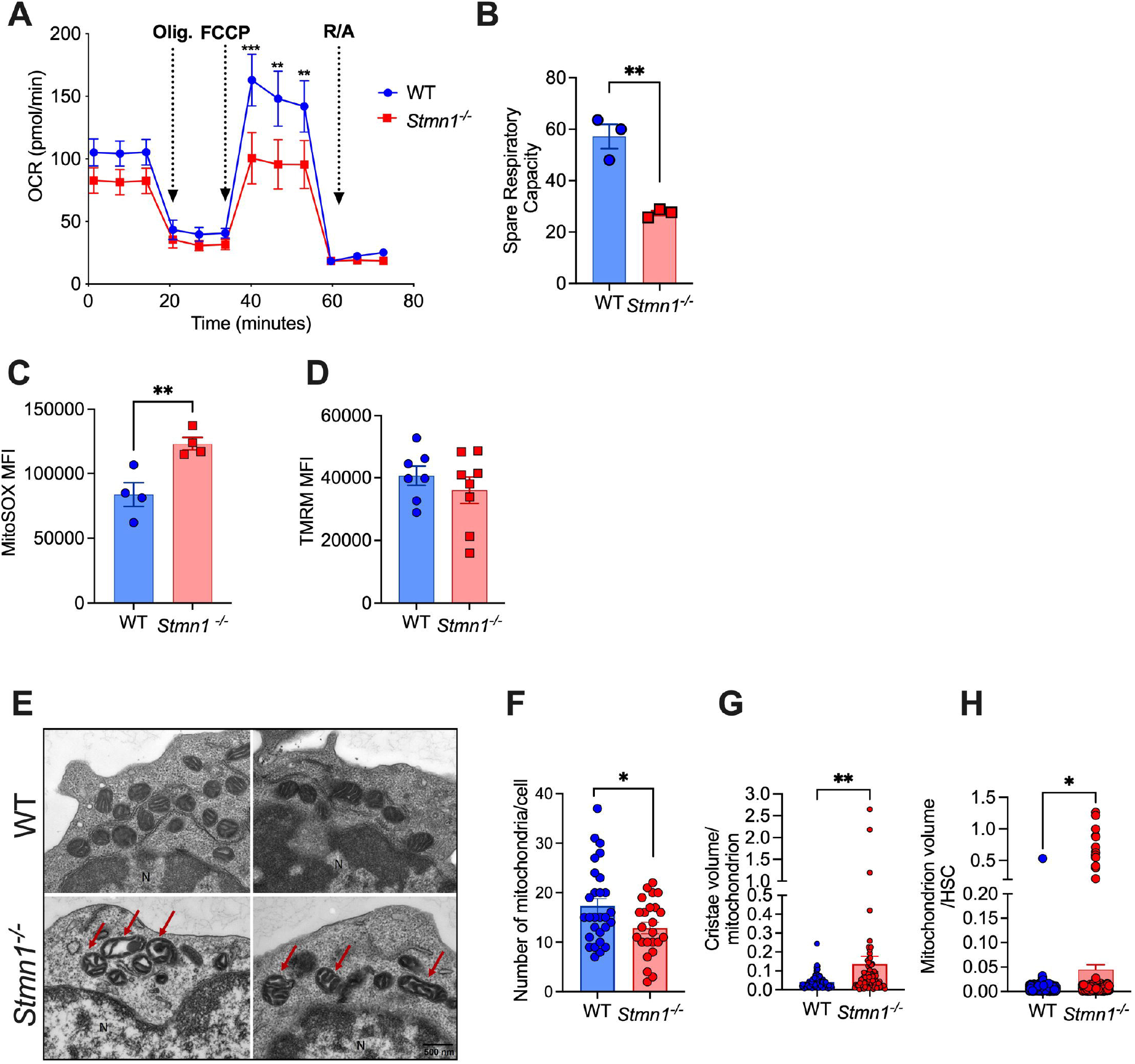
*Stmn1*^*-/-*^ HSPCs have decreased mitochondrial function. (A-B) Mitochondrial function (oxygen consumption rate, OCR) was measured on c-Kit+ bone marrow HSPCs from WT and *Stmn1*^*-/-*^ mice using the Seahorse analyzer. *Stmn1*^*-/-*^ HSPCs have impaired maximal respiratory capacity (shown following FCCP addition) compared to WT HSPCs (A) and reduced spare respiratory capacity (B, difference between maximal respiratory capacity and basal respiration prior to oligomycin addition). Data are representative of four biological replicates over three independent experiments. (C-D) Shown are HSC mitochondrial superoxide production measured using MitoSOX red (C) and mitochondrial membrane potential measured using TMRM (Tetramethylrhodamine methyl ester, (D) by flow cytometry. MFI= mean fluorescence intensity. (E) Representative Transmission Electron Microscopy micrographs (25000X magnification) of WT and *Stmn1*^*-/-*^ HSCs showing abnormal mitochondria morphology highlighted by red arrows. *Stmn1*^*-/-*^ HSCs have reduced mitochondria per cell (F), increased cristae volume per mitochondria (G), and increased mitochondria volume per HSC (H). Data represent the combination of two independent experiments. Error bars represent mean +/-SEM. * p<0.05, **p<0.01, ***p<0.001 by unpaired Student’s *t* test or (for G-H) Mann-Whitney.

Next, to examine mitochondrial morphology, we performed transmission electron microscopy (TEM) on isolated HSCs (Lineage-c-Kit+ Sca-1+ CD150+ CD48-cells) from WT and *Stmn1*^*-/-*^ mice. As shown in representative micrographs in Figure 5E, the mitochondria of *Stmn1*^*-/-*^ HSCs have markedly abnormal morphology. While overall mitochondria numbers per cell are decreased in *Stmn1*^*-/-*^ HSCs (Figure 5F), they have increased cristae volume and mitochondrial volume compared to WT (Figure 5G-H). To determine whether these mitochondrial abnormalities are unique to HSCs, we also sorted Gr-1+ myeloid cells from *Stmn1*^*-/-*^ and WT mice for TEM. As shown in Supplemental Figure 5A-B, the mitochondria in the *Stmn1*^*-/-*^ Gr-1+ cells also have some morphologic abnormalities, with reduced electron density and increased cristae volume, suggesting that Stathmin 1 may be important to maintain mitochondrial health beyond HSCs in more differentiated cell populations.

### HSC mitophagy is impaired in the absence of Stathmin 1

Together, our data suggest that Stathmin 1 is necessary to maintain normal mitochondrial and cellular function in HSCs. We considered that mitophagy, a specialized form of autophagy and a major regulator of mitochondrial quality, may be altered in *Stmn1*^*-/-*^ HSCs. High autophagic flux, and in particular the ability to clear damaged mitochondria by mitophagy, are key factors in maintaining HSC function.^23^ While the majority of autophagy-associated genes were not differentially expressed in *Stmn1*^*-/-*^ HSCs compared to WT (Supplemental Figure 6A-B), microtubules are still important for the trafficking of lysosomes needed for the completion of mitophagy. Thus, alterations in microtubule dynamics could potentially impair this process and lead to accumulation of dysfunctional mitochondria.^24-26^

**Figure 6.**
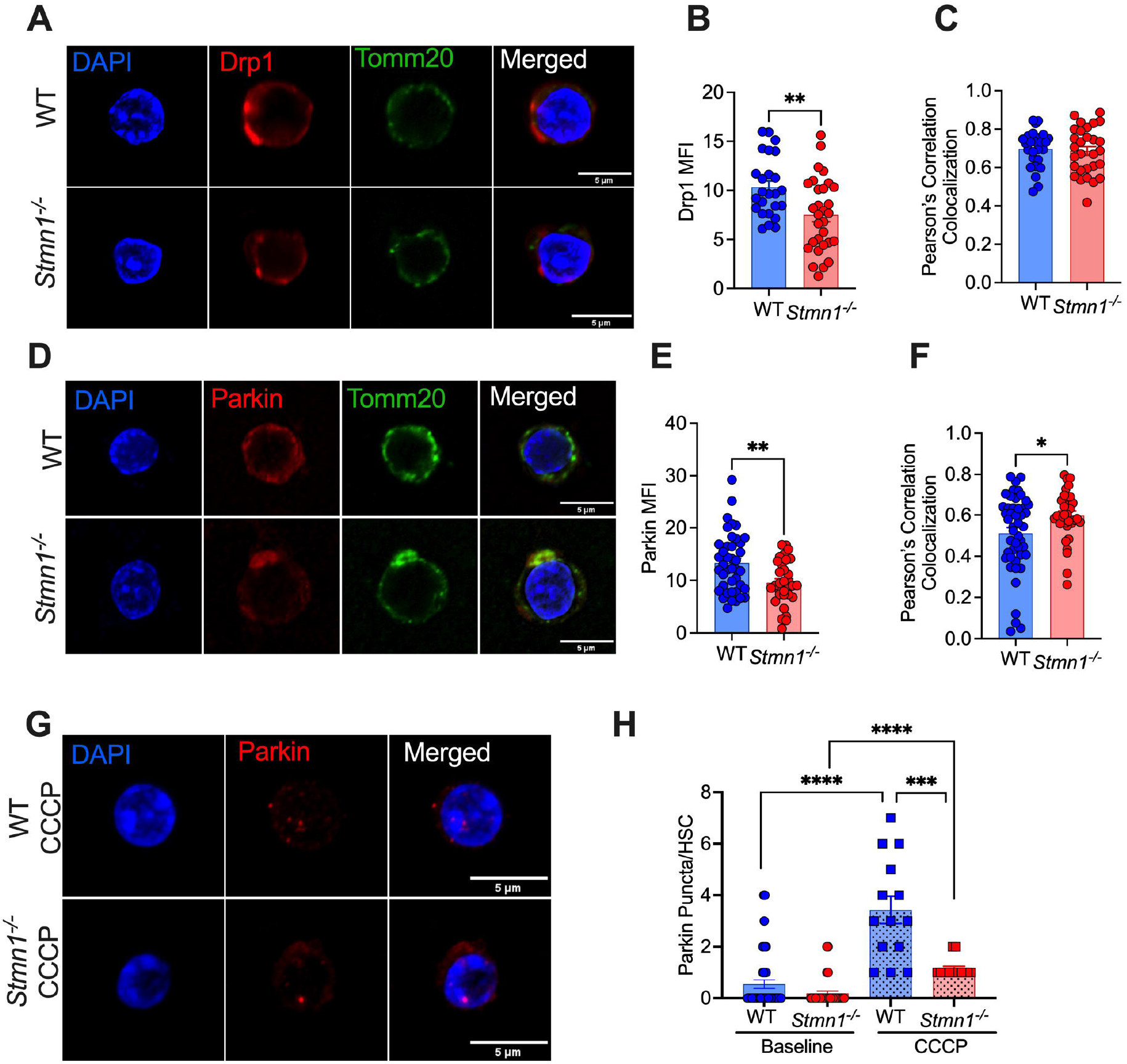
*Stmn1*^*-/-*^ HSCs have impaired mitophagy. (A) Representative confocal images of HSCs from WT and *Stmn1*^*-/-*^ mice stained for DAPI, Drp1 and Tomm20. (B-C) Quantification of Drp1 mean fluorescence intensity (MFI) and colocalization with the mitochondria marker Tomm20. (D) Representative confocal images of HSCs stained for DAPI, Parkin and Tomm20. (E-F) Quantification of Parkin MFI and colocalization with Tomm20. (G) Representative confocal images of HSCs stained for Parkin following treatment with CCCP. (H) Quantification of Parkin puncta per HSC at baseline and following CCCP treatment. Data represent the combination of 3 independent experiments. Error bars represent mean +/-SEM. *p<0.05, **<p0.01, ***p<0.001, ****p<0.0001 by unpaired Student’s t test.

To assess mitophagy, we first used confocal microscopy on sorted HSCs to evaluate the expression of Drp1 (Dynamin-related protein 1), a protein involved in mitochondrial fission that promotes mitophagy.^27,28^ While we did not detect any differences in localization of Drp1 to mitochondria (indicated by staining for the outer mitochondrial membrane protein Tomm20), Drp1 levels were decreased in *Stmn1*^*-/-*^ HSCs compared to WT (Figure 6A-C). Following Drp1-mediated fission, damaged and fragmented mitochondria are targeted for autophagic degradation by PTEN-induced kinase 1 (PINK1) and Parkin (an E3 ubiquitin ligase),^29^ so we next assessed the levels of Parkin in sorted HSCs. Parkin, which attaches to the outer membrane of damaged mitochondria,^30^ co-localized with mitochondria (Tomm20) in *Stmn1*^*-/-*^ HSCs. However, we detected overall reduced levels of Parkin in *Stmn1*^*-/-*^ HSCs compared to WT (Figure 6D-F), suggesting reduced mitophagy. Next, we treated the sorted HSCs with CCCP (carbonyl cyanide m-chlorophenyl hydrazone), an inhibitor of mitochondrial oxidative phosphorylation and inducer of PINK1/Parkin mediated mitophagy and stained for Parkin (Figure 6G). Compared to untreated baseline, CCCP treated WT HSCs had an expected increase in Parkin puncta, while *Stmn1*^*-/-*^ HSCs had a significantly diminished increase in puncta consistent with decreased mitophagy (Figure 6H).

Following Parkin-mediated ubiquitination, mitochondria fuse with LC3-containing autophagosomes.^31,32^ LC3 (microtubule-associated protein light chain 3) is thus a key protein in the autophagy pathway,^33^ and we next measured LC3 levels and colocalization with mitochondria (Tomm20) in sorted HSCs both at baseline and following treatment with an inhibitor of lysosomal proteolysis, leupeptin. Autophagic flux is measured by the accumulation of LC3 in the presence of leupeptin.^34^ At baseline, LC3 levels measured by immunofluorescence are similar between WT and *Stmn1*^*-/-*^ HSCs, as is the colocalization with Tomm20 (Figure 7A-B). However, following leupeptin treatment, LC3 levels were significantly higher in the WT HSCs compared to *Stmn1*^*-/-*^ HSCs (Figure 7C-D), indicating reduced autophagic flux in *Stmn1*^*-/-*^ HSCs compared to WT.

**Figure 7.**
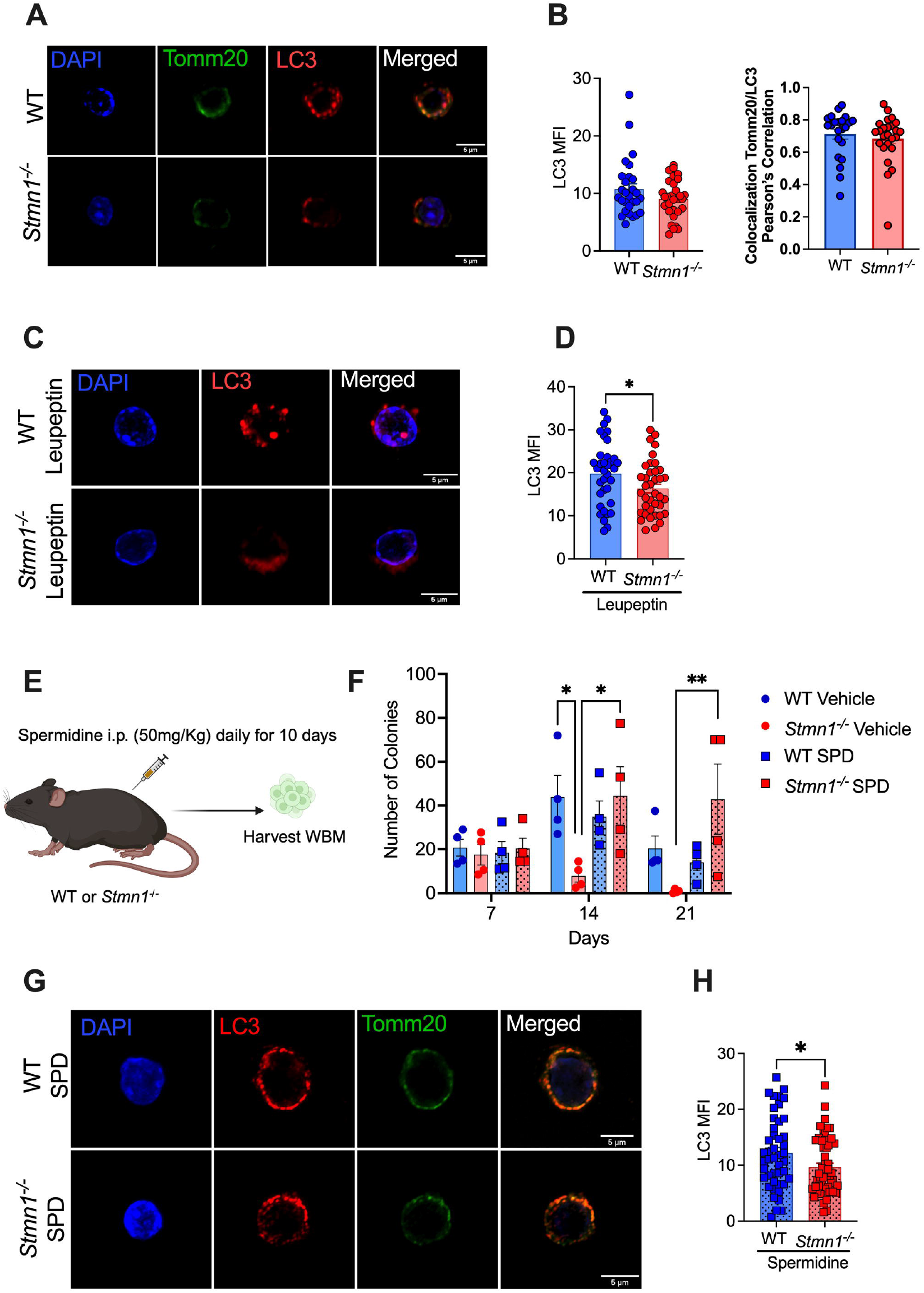
*Stmn1*^*-/-*^ HSCs have reduced autophagic flux. (A) Representative confocal images of HSCs from WT and *Stmn1*^*-/-*^ mice stained for Tomm20 and LC3. (B) Quantification of LC3 MFI at baseline and co-localization of LC3 and Tomm20. (C-D) Sorted HSCs were treated with leupeptin. Shown are representative confocal images of WT and *Stmn1*^*-/-*^ HSCs stained for LC3 following leupeptin treatment (C), and quantification of LC3 MFI in the treated cells (D). Data represent the combination of three independent experiments. (E) WT and *Stmn1*^*-/-*^ mice were treated with spermidine (50 mg/kg once daily) or vehicle for 10 days. (F) Bone marrow from spermidine treated and vehicle control mice was plated in MethoCult 3434 (1×10^4^ cells per well), and colonies were scored after 7 days. Cells were then replated (5×10^3^ cells/well) and scored every 7 days. Data represent 4 samples over 3 independent experiments. (G) Representative confocal images of HSCs stained for LC3 and Tomm20 from WT and *Stmn1*^*-/-*^ mice treated with spermidine. (H) Quantification of LC3 MFI in HSCs from spermidine treated mice. Data represent the combination of two independent experiments. Error bars represent mean +/-SEM. *p<0.05, **<p0.01 by unpaired Student’s t test or 2-way ANOVA.

### Autophagy stimulation improves Stmn1^-/-^ HSPC function

Together, these data show that Stathmin 1 is important for proper mitophagy and suggest that the reduced function of *Stmn1*^*-/-*^ HSCs may be due to impairment in this process. We therefore reasoned that stimulation of autophagy/mitophagy may improve *Stmn1*^*-/-*^ HSPC function, and to this end we treated mice with the autophagy inducer spermidine (50 mg/kg daily x 10 days, Figure 7E).^35^ As shown in Fig. 7F, WBM from spermidine treated *Stmn1*^*-/-*^ mice produced colonies with improved serial replating capacity in MethoCult compared to vehicle treated controls, suggesting some improvement in HSPC function with autophagy stimulation. Notably, while spermidine treatment increased LC3 levels in both WT and *Stmn1*^*-/-*^ HSCs (consistent with autophagy stimulation), the levels in *Stmn1*^*-/-*^ HSCs did not reach those of WT (Figure 7G-H), supporting the idea that autophagy is impaired in the absence of Stathmin 1.

## Discussion

HSCs display unique metabolic characteristics compared to differentiated cells. Specifically, they are characterized by high levels of relatively inactive mitochondria, and they rely heavily on glycolysis over oxidative phosphorylation (OXPHOS) for ATP production.^19,36,37^ Because of their low mitochondrial OXPHOS, they produce low levels of ROS. In addition, HSCs have high autophagic flux, and they rely on high levels of mitophagy to maintain mitochondrial health. Reduced autophagy and persistently elevated ROS are both associated with loss of HSC stemness and reduced function.^38,39^ HSCs are very sensitive to alterations in mitochondrial metabolism, and a switch from glycolysis to OXPHOS and increased ROS production are associated with HSC differentiation.^19^ Further, mutations that compromise mitochondrial function or increase ROS are associated with a reduction in the HSC pool and loss of HSC function,^19^ and alterations in mitochondrial function are thought to contribute to the decline in HSC function with normal aging. Aged HSCs display a loss of self-renewal and regenerative potential, and they are characterized by impaired mitophagy, mitochondria accumulation and dysfunction, and elevated ROS.^37^

Despite the importance of tight mitochondrial quality control to HSC function, the mechanisms that maintain this process in HSCs are not well understood. Herein, we identified Stathmin 1 as a critical regulator of HSC mitochondrial quality and cellular function in HSCs. Stathmin 1 has previously been implicated in erythro-megakaryopoiesis,^13^ however this is the first report to our knowledge of a role for Stathmin 1 in HSCs. Further, while a prior study found Stathmin 1 to promote mitochondrial integrity in stressed hepatocytes via down-regulation of c-Jun N-terminal kinase (JNK),^40^ and a second study showed that Stathmin 1 promotes paclitaxel-induced autophagy in osteosarcoma cell lines,^41^ this is the first report of a role for Stathmin 1 specifically in supporting mitophagy.

While our data clearly support a role for Stathmin 1 in regulating mitochondrial function in HSCs, several important questions remain. First, while mitochondrial quality control processes such as mitophagy and fission rely on proper microtubule dynamics, the specific role of Stathmin 1 is not clear. Indeed, microtubules have been shown to impair Drp1 assembly on mitochondria and thereby less mitochondria fragmentation.^28,42^ This fission process, followed by mitophagy, play a vital role in ensuring that damaged mitochondria are removed from the pool of healthy mitochondria. The exact role of Stathmin 1 in these processes is not known, much less the signaling system that coordinates the activity of Stathmin 1 with mitochondria fission and mitophagy. Further studies are needed to delineate the precise effects of Stathmin 1 loss on microtubules in HSCs, as well as to detail how these changes affect the association of microtubules with mitochondria, lysosomes, and other components of the mitophagy machinery. Next, it is not clear if Stathmin 1 is uniquely required to support mitophagy in HSPCs compared to other cell types. Stathmin 1 is highly expressed in HSPCs,^9^ with much lower levels in terminally differentiated effector blood cells.^10^ Further, while *Stmn1*^*-/-*^ mice have abnormalities in both erythroid and megakaryocyte lineage cells,^13^ they have surprisingly modest hematopoietic defects given their markedly impaired HSCs. Our TEM data, however, show mitochondrial abnormalities in *Stmn1*^*-/-*^ granulocytes, suggesting that Stathmin 1 may also be required to maintain mitochondrial quality control in more differentiated hematopoietic cells. Further studies are needed to confirm this hypothesis, and carefully elucidate whether mitochondrial metabolism and cellular function is altered throughout hematopoiesis in the absence of Stathmin 1.

Next, we have focused our current study to the role of Stathmin 1 in mitochondrial metabolism and mitophagy, and it is not clear whether it may be important in other autophagy-dependent processes. For example, HSCs rely heavily on the autophagy-lysosome system to degrade misfolded or damaged proteins, forming so-called aggresomes.^43^ These complexes of misfolded proteins traffic via microtubules to perinuclear locations where they are sequestered in a cage of vimentin-containing intermediate filaments and degraded by aggrephagy, a selective form of autophagy. Additional studies are needed to comprehensively interrogate the extent of autophagy dysregulation with Stathmin 1 loss.

Finally, the role of Stathmin 1 in malignant HSPCs is not clear. Stathmin 1 is highly expressed in both myeloid and lymphoid leukemias, where it has been shown to promote cell growth and survival.^5,6,8,10^ Leukemic cells are metabolically distinct from healthy HSPCs, with a stronger reliance on oxidative phosphorylation,^44,45^ suggesting that elevated levels of Stathmin 1 may facilitate leukemogenesis via promotion of mitochondrial activity. While inhibition of Stathmin 1 has been shown to induce cell cycle arrest and apoptosis in leukemia cell lines,^46,47^ the specific mechanisms by which it promotes leukemic cell growth and survival are not well understood. Further studies are thus needed to determine whether Stathmin 1 regulates mitochondrial quality in malignant HSPCs.

## Acknowledgments

The authors thank the DCM of Washington University for animal care. This work was supported by funding from The Children’s Discovery Institute of St. Louis Children’s Hospital Foundation (CJL, LGS) and the National Heart, Lung, and Blood Institute (R01 HL134896; LGS) and the contributions of D.J.K were supported by funding from the National Institute for General Medical Science (R01 GM136925).

## Author Contributions

L.C. and L.G.S. designed the research and wrote the manuscript; L.C., M.S., Q.D., M.W., L.V., A.J., P.H., C.P., J.W., N.R., Z.J.G., W.Y., and C.R.Z. performed and analyzed the results of the experiments; L.C. and M.S. prepared the figures; G.A.C., C.J.L., R.A.J.S., W.L.B., S.S., W.L. and D.J.K. provided critical resources and provided supervision of personnel.

## Declaration of Interests

The authors declare no competing interests.

## Notes

### Competing Interest Statement

The authors have declared no competing interest.

