## Supplemental Materials for "Stathmin 1 regulates mitophagy and cellular function in hematopoietic stem cells"

Chiquetto, L. et al

Supplemental Methods

Supplemental Figures 1-6

Supplemental Table 1. Antibodies and Other Reagents

Supplemental Table 2. Differentially expressed genes in *Stmn1*$^{-/-}$ vs WT HSCs

#### Supplemental Methods

*Cell sorting*
Bone marrow cells were obtained by crushing bones in FACS Buffer as previously described.[1] Cell suspensions were filtered using a 40-μm filter before staining with primary antibodies (Supplemental Table 1) on ice for 40 min. Cells were enriched for c-Kit using a biotin-conjugated CD117, followed by streptavidin-conjugated magnetic beads using MACS gravity positive labelling (Miltenyi Biotec, Auburn, CA, USA) before double sorting Lineage- c-Kit+ Sca-1+ CD150+ CD48- HSCs on a BD FACSAria Fusion (BD Biosciences, Franklin Lakes, NJ, USA). Non-viable cells were excluded by 4', 6-diamidino-2-phenylindole (DAPI) staining.

*Cell cycle analysis*
Bone marrow cells were stained with surface markers (Supplemental Table 1) and fixed with BD Cytofix/Cytoperm (BD Biosciences). Cells were then blocked with 5% goat serum and stained with mouse anti-human Ki67 (clone B56; BD Pharmingen) per manufacturer's instructions. Samples were washed and resuspended in DAPI- containing FACS buffer for analysis by flow cytometry. For G-CSF treatment, granulocyte-colony stimulation factor (G-CSF; Amgen, Thousand Oaks, CA, USA) was diluted in sterile phosphate buffered saline (PBS) and injected subcutaneously (3.75 μg/100 μL) every 12 hours for a total of 2 injections per mouse. Control mice were injected with vehicle (PBS). Bone marrow was analyzed as above after 2 hours following the last injection.

*Apoptosis analysis*
Bone marrow cells were isolated as described above, stained for surface markers, then washed with Annexin V binding buffer and staining for Annexin V using the Annexin V Apoptosis Detection Kit (eBioscience, San Diego, CA, USA) according to manufacturer instructions and resuspended in DAPI-containing FACs buffer for analysis by flow cytometry.

*Homing assay*
$1x10^5$ LSK (Lineage- cKit+ Sca-1+) cells from WT and *Stmn1*$^{-/-}$ mice (CD45.2) were sorted and transplanted retro-orbitally into lethally irradiated WT (CD45.1) mice. After 18 hours, bone marrow was harvested from recipient mice and bone marrow chimerism was analyzed by flow cytometry.

*5-FU recovery analysis*
Mice were weighed, peripheral blood was collected, and 5-fluorouracil (5-FU) was injected intraperitoneally (IP; 100 mg/kg) on day zero. Peripheral blood was collected every 2-3 days and complete blood counts were analyzed on the Element HT5 automated cell counter until blood counts returned to baseline levels.

*MethoCult colony growth assays*
Whole bone marrow cells ($1x10^4$ cells) or sorted HSCs mice were incubated in MethoCult GF M3434 (Stem Cell Technologies, Vancouver, BC, Canada). Colonies were scored after 7 days at 37°C with 5% $CO_2$. For serial replating analyses, wells were washed and replated at a density of $5x10^3$ cells per well every 7 days.

*Spermidine treatment*

Spermidine (Sigma-Aldrich, St. Louis, MO, USA) was injected IP (50 mg/kg) daily for 10 days. 16 hours following the last treatment, bone marrow was harvested for MethoCult growth assays and HSCs were sorted for LC3 analysis by confocal microscopy and Western blot.

*Mitochondria membrane potential and mitochondrial ROS measurements*

To measure mitochondrial membrane potential, whole bone marrow was stained with HSC markers. After washing, cells were stained with 50 nM tetramethylrhodamine (TMRM) dye at 37°C for 30 minutes. CCCP 5uM (Invitrogen, Waltham, MA, USA) was used as a control, incubated at 37°C for 5 minutes. The TMRM signal was measured on a Cytek Aurora flow cytometer. For measuring mitochondrial superoxide production, whole bone marrow was stained with MitoSOX Red (Invitrogen) for 30 minutes at 37°C. Mitochondria ROS levels were obtained by Cytek Aurora flow cytometer using the MitoSOX channel. Mean fluorescence intensity (MFI) was calculated using FlowJo software.

*RNA Sequencing*

HSCs were double sorted directly into lysis buffer. RNA was isolated using the NucleoSpin RNA XS Kit (Machery-Nagel, Bethlehem, PA, USA). Samples were then prepared according to library kit manufacturer's protocol, indexed, pooled, and sequenced on an Illumina NovaSeq 6000. Base calls and demultiplexing were performed with Illumina's bcl2fastq2 software. RNA-seq reads were aligned and quantitated to the Ensembl release 101 primary assembly with an Illumina DRAGEN Bio-IT server running version 3.9.3-8 software. All gene counts were then imported into the R/Bioconductor package EdgeR1 and TMM normalization size factors were calculated to adjust for differences in library size. Differential expression analysis results were filtered for genes with Benjamini-Hochberg false-discovery rate adjusted p-values less than or equal to 0.05. Global perturbations in known Gene Ontology (GO) terms, MSigDb, and KEGG pathways were detected using the R/Bioconductor package GAGE4. The R/Bioconductor package heatmap35 was used to display heatmaps across groups of samples for autophagy-related genes.

*Microtubule network staining*

For microtubule network staining, cells were sorted into StemSpan SFEM media with TPO (100ng/mL) and SCF (100ng/mL) and incubated for 24 hours at 37C and 5% C02. Cells were then plated in poly-L-lysine coated glass coverslips, rested for 45 minutes at 37°C, washed with warm PBS, fixed with ice-cold methanol for 10 minutes at -20°C and washed with ice-cold PBS. They were then permeabilized in 0.5% v/v TritonX-100 (Pierce Biotechnology, Waltham, MA, USA) in PBS for 10 minutes before being blocked in 3% BSA in PBS with 0.05% Tween-20 (Millipore Sigma, Burlington, MA, USA). Primary antibodies were incubated in the block buffer for 1 hour at RT before washing twice with 0.01% v/v TritonX-100 in PBS. Secondary antibodies and dyes were also incubated in the block buffer for one hour at RT before being washed twice with 0.01% v/v TritonX-100 in PBS. Finally, coverslips were incubated with 0.25 µg/ml of DAPI for 10 minutes at room temperature before being washed twice with 0.01% v/v TritonX-100 in PBS and mounted in a solution of ProLong Glass Antifade mountant (Invitrogen) and allowed to cure for 24 h before imaging. Images were captured using a Nikon AXR confocal system on an Eclipse Ti2 inverted microscope equipped with a Nikon Spatial Array Confocal (NSPARC) detector using a 60× (1.42 NA) Plan-Apo oil immersion objective. Image stacks were captured using galvanometer (resonance) scanning at 16-bit with a resolution of 2048x2048, axial spacing of 0.2 µm, and a pinhole size of 2 using the Nikon Elements Software package. Image analysis was performed using the software Fiji.

*Transmission Electron Microscopy*
For ultrastructural analyses, $2x10^4$ sorted HSCs were fixed in 2% paraformaldehyde/2.5% glutaraldehyde (Ted Pella Inc., Redding, CA, USA) in 100 mM sodium cacodylate buffer for 2 hours at room temperature. Samples were washed in sodium cacodylate buffer and post-fixed in 1% osmium tetroxide (Ted Pella Inc.) for 1 hour at room temperature. Samples were then rinsed extensively in $dH_20$ prior to en bloc staining with 1% aqueous uranyl acetate (Electron Microscopy Sciences, Hatfield, PA, USA) for 1 hour. Following several rinses in $dH_20$, samples were dehydrated in a graded series of ethanol, and embedded in Eponate 12 resin (Ted Pella Inc). Ultrathin sections of 95 nm were cut with a Leica Ultracut UCT ultramicrotome (Leica Microsystems Inc., Bannockburn, IL), stained with uranyl acetate and lead citrate, and viewed on a JEOL 1200 EX transmission electron microscope (JEOL USA Inc., Peabody, MA) equipped with an AMT 8 megapixel digital camera and AMT Image Capture Engine V602 software (Advanced Microscopy Techniques, Woburn, MA). Measurements of mitochondria cristae were done on ImageJ following the protocol previously described.[2]

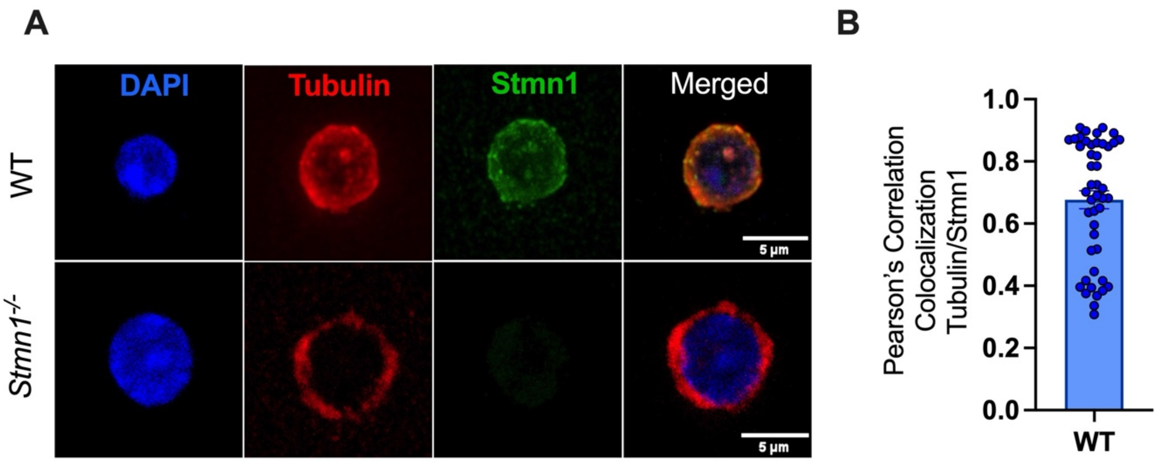

**Supplemental Figure 1. Stathmin 1 is expressed in HSCs and co-localizes with tubulin. (A)** Shown are representative confocal immunofluorescence images of sorted HSCs (Lin- c-Kit+ Sca-1+ CD150+ CD48- cells) from WT and *Stmn1*$^{-/-}$ mice including DAPI (blue), alpha-tubulin (red), and Stathmin 1 (green), and colocalization of tubulin and Stathmin 1 by Pearson's coefficient in WT cells **(B)**.

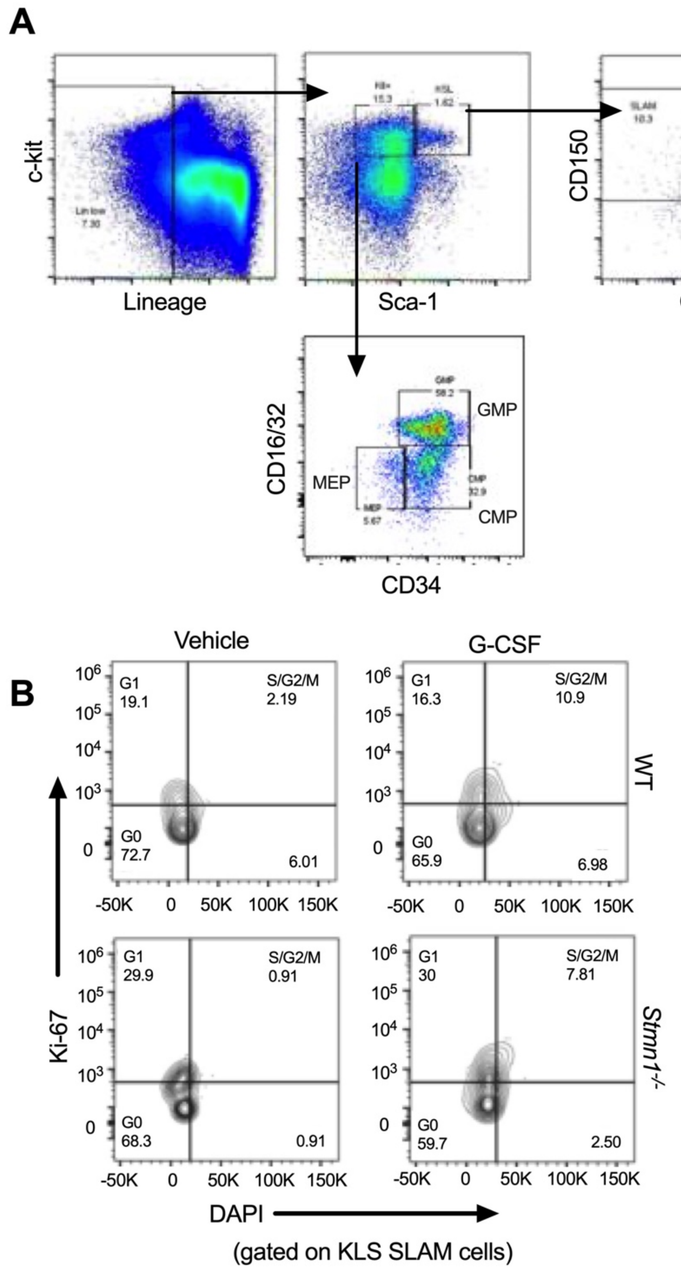

**Supplemental Figure 2. HSC and Ki67/DAPI gating strategies. (A)** Representative flow plots of hematopoietic stem cell and myeloid progenitor gating strategy (related to main Figure 1E-H). **(B)** Representative flow plots of HSCs further gated for Ki-67 and DAPI from WT and *Stmn1*$^{-/-}$ young adult mice treated +/- G-CSF (related to main Figure 3G-I).

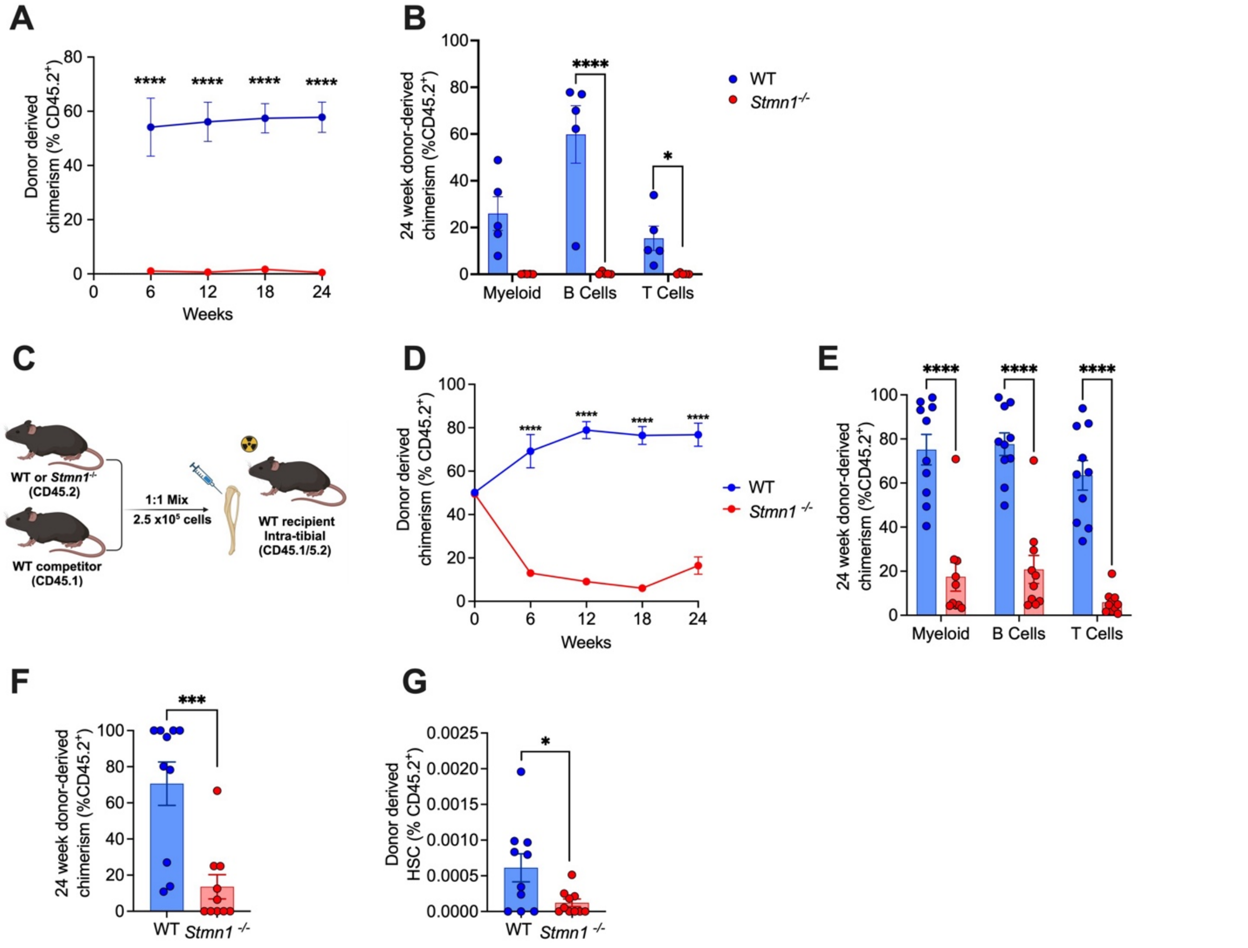

**Supplemental Figure 3. *Stmn1$^{-/-}$* HSCs have markedly impaired function. (A-B)** WBM from primary transplant recipient mice as described in main Figure 2A-C was harvested after 24 weeks and transplanted into new lethally irradiated recipients. Chimerism was assessed by peripheral blood (PB) flow cytometry every 6 weeks. Shown are total %CD45.2 donor derived cells in the PB over time **(A)**, and tri-lineage engraftment in the PB at 24 weeks **(B)**. n= 5 mice per group. **(C)** Outline of intra-tibial competitive whole BM transplants. WBM from WT or *Stmn1$^{-/-}$* mice (CD45.2) was mixed 1:1 with WBM from a WT competitor (CD45.1) and 2.5 x $10^5$ total cells were injected intra-tibially into lethally irradiated WT (CD45.1/CD45.2) recipients. Chimerism was assessed by PB flow cytometry every 6 weeks. Shown are total %CD45.2 donor derived cells in the PB over time **(D)**, tri-lineage engraftment in the PB at 24 weeks **(E)**, and %CD45.2 donor derived total cells **(F)** and HSCs **(G)** in the bone marrow at 24 weeks. n = 10 recipients per group over 2 independent transplants. Error bars represent mean +/- SEM. *p<0.05, ***p<0.001, ****p<0.0001 by unpaired Student's t- test.

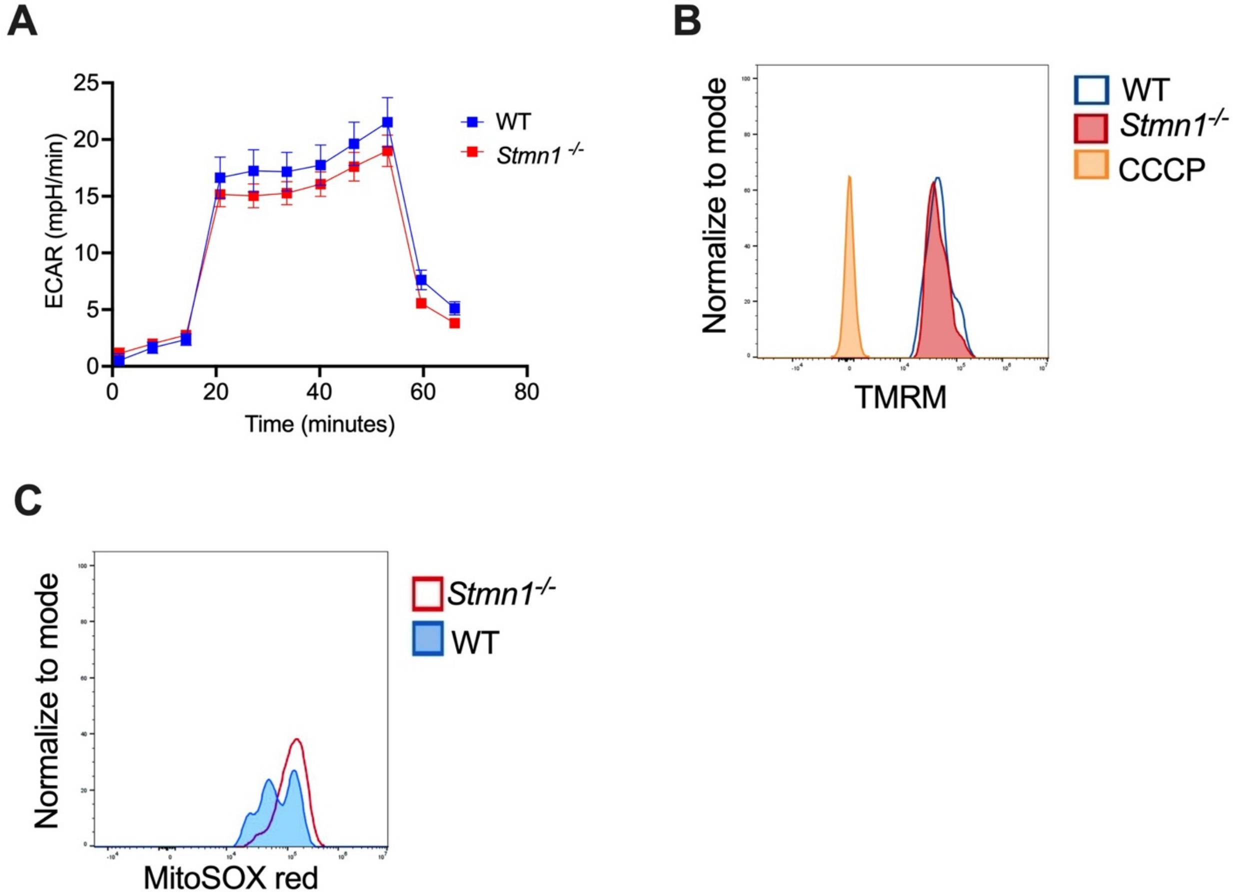

**Supplemental Figure 4. Glycolytic capacity is similar between *Stmn1*$^{-/-}$ and WT HSPCs. (A)** Glycolytic capacity of c-Kit+ HSPCs from WT and *Stmn1*$^{-/-}$ mice were measured using the Seahorse Glycolysis Stress test. Shown are combined data from two independent experiments. **(B)** Representative histogram of HSC mitochondria membrane potential determined by flow cytometry using the tetramethylrhodamine (TMRM) dye. CCCP 5uM was used as a control (related to main Figure 5D). **(C)** Representative histogram of MitoSOX Red from WT and *Stmn1*$^{-/-}$ HSCs by flow cytometry (related to main Figure 5C).

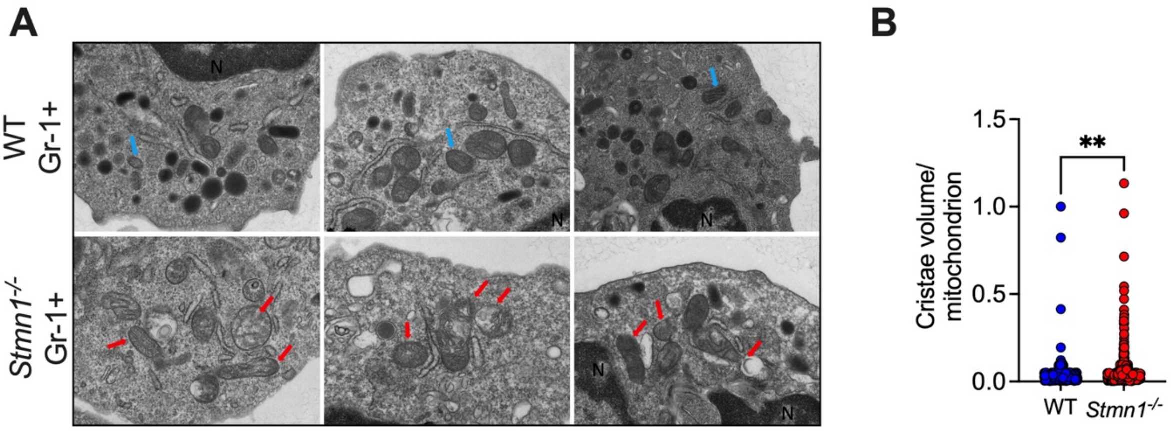

**Supplemental Figure 5. Mitochondrial abnormalities are also found in Gr-1+ myeloid cells from *Stmn1*$^{-/-}$ mice. (A)** Shown are representative Transmission Electron Microscopy (TEM) micrographs highlighting the mitochondria morphology in sorted Gr-1+ cells from WT (top row; blue arrows) and *Stmn1*$^{-/-}$ mice (bottom row; red arrows). **(B)** Quantification of cristae volume. **p<0.01 by Mann-Whitney. Shown are combined data from two independent experiments.

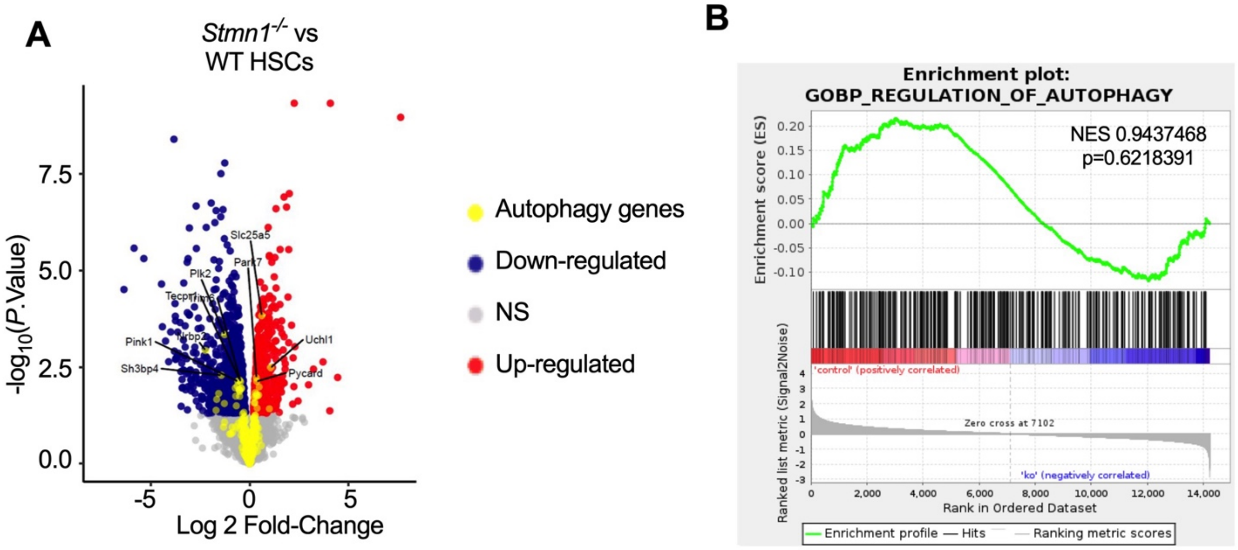

**Supplemental Figure 6. Most autophagy genes are not differentially expressed in *Stmn1*$^{-/-}$ HSCs. (A)** Volcano plot of differentially expressed genes between *Stmn1*$^{-/-}$ and WT HSCs. Autophagy- related genes are highlighted in yellow. **(B)** Enrichment plot for autophagy pathway genes comparing *Stmn1*$^{-/-}$ to WT HSCs.

Supplemental Table 1. Antibodies and Other Reagents

| Antibodies | Source | Identifier |
|---|---|---|
| Anti-mouse CD150 PE | Biolegend | CAT #115912 |
| Anti-mouse CD48 PE-Cy7 | eBiosciences | CAT #25-0481-80 |
| Anti-mouse CD48 APC | eBiosciences | CAT #17-0481-82 |
| Anti-mouse Flk2 APC | eBiosciences | CAT #17-1351-82 |
| Anti-mouse CD117 (c-Kit) APC-eFluor 780 | eBiosciences | CAT #47-1172-82 |
| Anti-mouse CD34 FITC | eBiosciences | CAT #11-0341-85 |
| Anti-mouse Sca-1 PerCP-Cy5.5 | eBiosciences | CAT #368-5981-80 |
| Anti-mouse CD16/32 e450 | eBiosciences | CAT #48-0161-82 |
| Anti-mouse Steptavidin PE-eFluor 610 | eBiosciences | CAT #61-4317-82 |
| Anti-mouse Steptavidin PE-Cy7 | eBiosciences | CAT# 25-4317-82 |
| Anti-mouse c-Kit Biotin | eBiosciences | CAT #13-1171-82 |
| Anti-mouse B220 Biotin | eBiosciences | CAT #13-0452-85 |
| Anti-mouse CD3e Biotin | eBiosciences | CAT #13-0031-82 |
| Anti-mouse Ter-119 Biotin | eBiosciences | CAT #13-5921-82 |
| Anti-mouse Gr-1 Biotin | eBiosciences | CAT #13-5931-82 |
| Anti-mouse B220 PE | eBiosciences | CAT 12-0453-82 |
| Anti-mouse CD3e PE | eBiosciences | CAT #12-0031-82 |
| Anti-mouse Ter-119 PE | eBiosciences | CAT #12-5921-82 |
| Anti-mouse Gr-1 PE | eBiosciences | CAT #12-5931-82 |
| Anti-mouse B220 FITC | eBiosciences | CAT #11-0452-82 |
| Anti-mouse CD3e FITC | eBiosciences | CAT #11-0031-82 |
| Anti-mouse Ter-119 FITC | eBiosciences | CAT #11-5921-82 |
| Anti-mouse Gr-1 FITC | Biolegend | CAT #108406 |
| Anti-mouse CD45.2 FITC | eBiosciences | CAT #11-0454-82 |
| Anti-mouse CD3e PerCP-Cy5.5 | eBiosciences | CAT #45-0031-82 |
| Anti-mouse CD45.1 APC | eBiosciences | CAT #17-0453-82 |
| Anti-mouse Gr-1 APC-eFluor 780 | eBiosciences | CAT #47-5931-82 |
| Anti-mouse Gr-1 PE-Cy7 | eBiosciences | CAT #25-5931-81 |
| Anti-mouse Annexin V APC | eBiosciences | CAT# 17-8007-74 |
| Anti-mouse Ki67 FITC | Biolegend | CAT #151212 |
| Stmn1 Rabbit Polyclonal | Proteintech | CAT #11157-1-AP |
| LC3B Rabbit Polyclonal | Invitrogen | CAT #PA5-32254 |
| Parkin Rabbit polyclonal | Invitrogen | CAT #PA5-13399 |
| DRP1 (DNM1L) Mouse Monoclonal | Invitrogen | CAT# MA5-26255 |
| Tom20 (F-10) | Santa Cruz Biotechnology | CAT #sc17764 |
| Tom20 (D8T4N) | Cellsignaling Technologies | CAT #72610S |
| anti-a-Tubulin (DM1A) | Milipore Sigma | CAT #T9026 |
| Alexa Fluor 594 Anti-Tubulin-alpha Mouse | Biolegend | CAT #627910 |
| Alexa Fluor 568 Donkey anti-Rabbit | Invitrogen | CAT #A10042 |
| Alexa Fluor 488 Goat anti-rabbit | Invitrogen | CAT #A32731 |
| Alexa Fluor 488 Goat anti-mouse | Invitrogen | CAT #A11001 |
| **Chemicals /Recombinant Proteins** | **Source** | **Catalog Number** |
| Methocult M3434 | Stemcell Technologies | CAT #03434 |
| StemSpan SFEM | Stemcell Technologies | CAT #09600 |
| TPO | Sigma-Aldrich | CAT #415952 |
| SCF | Sigma-Aldrich | CAT #SRP3234 |
| G-CSF | ThermoFisher | CAT #315-03 |
| Mitotempo | Sigma-Aldrich | CAT #SML0737 |
| Spermidine | Sigma-Aldrich | CAT #S0266 |
| Leupeptin | ThermoFisher Scientific | CAT #78435 |
| BD Cytofix/Cytoperm | BD Biosciences | CAT #554722 |
| BD Perm/Wash | BD Biosciences | CAT #554723 |
| Annexin V Binding Buffer 10X | Invitrogen | CAT #00-0055-56 |
| MitoProbe TMRM kit | Invitrogen | CAT #M20036 |
| MitoSOX Red kit | Invitrogen | CAT #M36008 |

### Supplemental Table 2. Differentially expressed genes in *Stmn1*$^{-/-}$ vs WT HSCs

| Feature_ID | external_gene_name | logFC | CI.L | CI.R | AveExpr | t | P.Value | adj.P.Val | B | linearFC | sample.ko5 | sample.ko4 | sample.ko3 | sample.ko2 | sample.ko1 | sample.wt5 | sample.wt4 | sample.wt3 | sample.wt2 | sample.wt1 |
|---|---|---|---|---|---|---|---|---|---|---|---|---|---|---|---|---|---|---|---|---|
| ENSMUSG00000 | Gm10036 | 4.08E+00 | 3.39E+00 | 4.77E+00 | 3.180833 | 1.24E+01 | 4.74E-10 | 3.32E-06 | 12.40647 | 16.90418 | 4.798918431 | 4.82465511 | 5.84765914 | 5.150549482 | 4.91043252 | 1.73684548 | 0.906823109 | 0.36384405 | 2.167975466 | 1.10062321 |
| ENSMUSG00000 | Gm10282 | 2.25E+00 | 1.87E+00 | 2.64E+00 | 6.403425 | 1.24E+01 | 4.72E-10 | 3.32E-06 | 13.36993 | 4.7711 | 8.020511925 | 6.82928207 | 7.554847311 | 7.94618523 | 7.39654991 | 5.48766528 | 5.512240199 | 5.29848121 | 4.63409893 | 5.35438887 |
| ENSMUSG00000 | Gm6594 | 7.63E+00 | 6.26E+00 | 9.00E+00 | -2.08136 | 1.17E+01 | 1.09E-09 | 5.09E-06 | 7.480898 | 198.3961 | 1.277714765 | 0.83183667 | 3.244222457 | 1.922291866 | 1.18104763 | -5.83861813 | -6.012516974 | -5.98736022 | -5.53010407 | -5.90207931 |
| ENSMUSG00000 | Fscn1 | -3.83E+00 | -4.57E+00 | -3.08E+00 | 3.199571 | -1.08E+01 | 4.02E-09 | 1.41E-05 | 10.04485 | -14.1807 | 0.748138506 | 1.6540833 | 1.449748827 | 1.445788954 | 0.80013851 | 5.44389117 | 5.028457625 | 4.86048013 | 5.473413841 | 5.09156675 |
| ENSMUSG00000 | Art4 | -1.26E+00 | -1.53E+00 | -9.88E-01 | 6.440894 | -9.81E+00 | 1.66E-08 | 4.65E-05 | 9.823734 | -2.39154 | 5.546434925 | 6.14579263 | 6.065420532 | 5.562263817 | 5.77514728 | 7.07931331 | 7.005161403 | 6.93355084 | 7.112624573 | 7.18322716 |
| ENSMUSG00000 | Ndn | -1.45E+00 | -1.78E+00 | -1.13E+00 | 5.580836 | -9.40E+00 | 3.13E-08 | 7.32E-05 | 9.235096 | -2.73401 | 4.680581589 | 4.99975208 | 4.60704622 | 4.505512732 | 5.60888532 | 6.13973482 | 6.198458704 | 6.14224479 | 6.613598006 | 6.31254338 |
| ENSMUSG00000 | Selp | 2.01E+00 | 1.52E+00 | 2.49E+00 | 5.928981 | 8.66E+00 | 1.03E-07 | 2.05E-04 | 8.002421 | 4.015091 | 6.377320844 | 7.20797273 | 6.595687297 | 7.388937879 | 6.44059582 | 5.19750052 | 4.442728039 | 4.61748948 | 5.513292386 | 5.50828724 |
| ENSMUSG00000 | Sell | 1.74E+00 | 1.31E+00 | 2.16E+00 | 6.24952 | 8.54E+00 | 1.26E-07 | 2.21E-04 | 7.762577 | 3.330846 | 7.404581799 | 7.04161771 | 6.950375942 | 7.477705804 | 6.34696346 | 5.83513261 | 4.968336633 | 5.35749046 | 5.134343214 | 5.97865212 |
| ENSMUSG00000 | Pde2a | -1.94E+00 | -2.43E+00 | -1.45E+00 | 5.02294 | -8.32E+00 | 1.80E-07 | 2.81E-04 | 7.53265 | -3.84229 | 3.269759181 | 4.78895252 | 4.12569756 | 3.591290481 | 5.03016977 | 6.12538332 | 5.833145563 | 6.07321413 | 5.608679686 | 5.78310749 |
| ENSMUSG00000 | Hk3 | 1.33E+00 | 9.86E-01 | 1.68E+00 | 6.003568 | 8.12E+00 | 2.53E-07 | 2.84E-04 | 7.077412 | 2.516078 | 6.701614968 | 6.93716127 | 5.971669958 | 6.92466767 | 6.37286434 | 5.22275306 | 5.19388694 | 5.49997739 | 5.945122707 | 5.26596596 |
| ENSMUSG00000 | Stmn1 | -1.35E+00 | -1.70E+00 | -9.97E-01 | 9.085948 | -8.09E+00 | 2.67E-07 | 2.84E-04 | 6.915655 | -2.54427 | 8.811725952 | 7.91060152 | 8.591821838 | 8.75358306 | 8.16774097 | 10.0294033 | 10.03872809 | 9.80497001 | 8.912212255 | 9.83869492 |
| ENSMUSG00000 | Tns2 | -2.70E+00 | -3.40E+00 | -2.01E+00 | 4.128156 | -8.22E+00 | 2.15E-07 | 2.84E-04 | 7.288852 | -6.5203 | 1.869324852 | 3.70024075 | 2.735392804 | 2.061557501 | 3.61490138 | 5.60173199 | 5.364151088 | 5.12963345 | 6.133009237 | 5.07161806 |
| ENSMUSG00000 | Cd53 | 1.86E+00 | 1.38E+00 | 2.34E+00 | 5.346511 | 8.18E+00 | 2.29E-07 | 2.84E-04 | 7.23214 | 3.642281 | 6.42696549 | 5.60263387 | 6.193417258 | 6.781404225 | 5.95737163 | 4.95985367 | 3.901868159 | 4.34019242 | 4.531942067 | 4.76946176 |
| ENSMUSG00000 | Obscn | -1.70E+00 | -2.14E+00 | -1.25E+00 | 5.319565 | -8.06E+00 | 2.83E-07 | 2.84E-04 | 7.070911 | -3.23846 | 3.901442413 | 5.0801976 | 4.033138907 | 4.10984609 | 5.43669781 | 6.16384531 | 6.075277331 | 6.0937903 | 6.504351024 | 5.79705931 |
| ENSMUSG00000 | Mmrn1 | -1.46E+00 | -1.85E+00 | -1.07E+00 | 7.881127 | -7.85E+00 | 4.10E-07 | 3.83E-04 | 6.511782 | -2.75072 | 6.831000842 | 7.99935428 | 7.188304752 | 6.531724649 | 7.66069425 | 8.4524806 | 8.539371493 | 8.72886978 | 8.385681963 | 8.49378949 |
| ENSMUSG00000 | Sult1a1 | -1.76E+00 | -2.24E+00 | -1.27E+00 | 5.197546 | -7.65E+00 | 5.79E-07 | 5.07E-04 | 6.366775 | -3.37689 | 3.384239855 | 4.95447629 | 4.199388251 | 4.350602234 | 4.94015068 | 6.33908792 | 5.812036101 | 6.20143877 | 5.911895975 | 5.88214655 |
| ENSMUSG00000 | Arhgef28 | -3.05E+00 | -3.90E+00 | -2.19E+00 | 1.971747 | -7.47E+00 | 7.95E-07 | 5.86E-04 | 5.452728 | -8.25469 | 0.862471181 | 1.20191156 | -0.31578592 | 1.269362387 | -1.35042117 | 3.76315265 | 3.81879027 | 3.75579117 | 3.254530775 | 3.45767025 |
| ENSMUSG00000 | Padi4 | 9.36E-01 | 6.73E-01 | 1.20E+00 | 7.590504 | 7.49E+00 | 7.70E-07 | 5.86E-04 | 5.859388 | 1.913403 | 7.960703833 | 8.46599432 | 7.819552849 | 8.054874663 | 7.9474122 | 7.40856767 | 6.949836835 | 7.1433712 | 7.120723303 | 7.03400647 |
| ENSMUSG00000 | Igf2bp2 | -2.19E+00 | -2.81E+00 | -1.58E+00 | 3.208197 | -7.49E+00 | 7.68E-07 | 5.86E-04 | 6.035671 | -4.57649 | 1.424350069 | 1.60137625 | 3.065805096 | 2.6260559 | 1.9772662 | 4.36961622 | 4.376576548 | 4.55856955 | 3.991496369 | 4.09085903 |
| ENSMUSG00000 | Dpp4 | -1.28E+00 | -1.66E+00 | -8.99E-01 | 4.474195 | -7.12E+00 | 1.51E-06 | 1.06E-03 | 5.453755 | -2.42372 | 3.976362148 | 3.73582645 | 3.193093416 | 4.053066041 | 4.41192765 | 5.26751291 | 5.138715256 | 5.06748825 | 4.804169214 | 5.09378675 |
| ENSMUSG00000 | Apbb1 | -1.13E+00 | -1.48E+00 | -7.87E-01 | 4.830286 | -6.92E+00 | 2.20E-06 | 1.47E-03 | 5.056829 | -2.19153 | 3.969999645 | 4.55795164 | 4.368534133 | 3.917434931 | 4.75810329 | 5.44157266 | 5.441268082 | 5.24085722 | 5.348713214 | 5.25842244 |
| ENSMUSG00000 | 6330403K07Rik | -5.84E+00 | -7.65E+00 | -4.04E+00 | -0.71564 | -6.82E+00 | 2.65E-06 | 1.61E-03 | 2.33149 | -57.3537 | -4.26681184 | -1.50296041 | -5.807639016 | -5.67315212 | 1.03914564 | 1.61259298 | 1.708582215 | 1.6492644 | 1.904524157 | 2.18006973 |
| ENSMUSG00000 | Lat2 | 1.54E+00 | 1.06E+00 | 2.02E+00 | 3.444053 | 6.78E+00 | 2.87E-06 | 1.61E-03 | 4.856706 | 2.905563 | 4.477663245 | 3.95234709 | 4.803902255 | 4.059931013 | 3.62737162 | 3.03703908 | 2.317772509 | 2.83766214 | 2.631985091 | 2.69485583 |
| ENSMUSG00000 | Gimap4 | 1.97E+00 | 1.36E+00 | 2.59E+00 | 4.152187 | 6.78E+00 | 2.88E-06 | 1.61E-03 | 4.818282 | 3.928235 | 4.664910613 | 5.12522993 | 5.122359046 | 5.358204477 | 5.40820181 | 1.76871218 | 3.796447201 | 3.22452807 | 3.676910249 | 3.37637015 |
| ENSMUSG00000 | Gm27022 | -2.69E+00 | -3.53E+00 | -1.86E+00 | 2.122957 | -6.81E+00 | 2.70E-06 | 1.61E-03 | 4.449844 | -6.46742 | 0.473396176 | 0.42849573 | 0.586051748 | 0.719165304 | 1.52016056 | 3.40650669 | 3.447037597 | 3.238522 | 4.399479832 | 3.01075016 |
| ENSMUSG00000 | Mpl | -9.85E-01 | -1.29E+00 | -6.77E-01 | 10.06902 | -6.73E+00 | 3.13E-06 | 1.69E-03 | 4.39249 | -1.97911 | 9.221572287 | 10.1943373 | 9.53349754 | 9.1603804 | 9.9673306 | 10.4291378 | 10.45437202 | 10.5483252 | 10.74327267 | 10.4379968 |
| ENSMUSG00000 | Fbp1 | 9.96E-01 | 6.78E-01 | 1.31E+00 | 5.025128 | 6.59E+00 | 4.12E-06 | 2.14E-03 | 4.35899 | 1.994404 | 5.496400531 | 5.48208456 | 5.005340455 | 5.78166123 | 5.74085553 | 4.6681853 | 4.38942915 | 4.41245174 | 4.600466492 | 4.67440504 |
| ENSMUSG00000 | Socs2 | 9.93E-01 | 6.74E-01 | 1.31E+00 | 6.275646 | 6.56E+00 | 4.35E-06 | 2.18E-03 | 4.170382 | 1.990895 | 6.884314667 | 6.40480997 | 6.632534584 | 6.779073675 | 7.23664373 | 6.07987246 | 5.390761402 | 5.83799612 | 5.678398557 | 5.83205447 |
| ENSMUSG00000 | Gem | 1.28E+00 | 8.66E-01 | 1.69E+00 | 4.820832 | 6.53E+00 | 4.63E-06 | 2.23E-03 | 4.267998 | 2.426146 | 5.779857469 | 5.72239165 | 5.38896311 | 5.072522205 | 5.46771359 | 4.37933956 | 3.783522635 | 4.64654319 | 3.991496369 | 3.9759716 |
| ENSMUSG00000 | Mettl24 | -5.35E+00 | -7.08E+00 | -3.61E+00 | -2.20051 | -6.50E+00 | 4.93E-06 | 2.24E-03 | 1.868663 | -40.7273 | -5.85177434 | -1.19262029 | -5.807639016 | -5.67315212 | -5.99427736 | 0.79073849 | 0.137230146 | 0.52043442 | 0.94562936 | 0.1202885 |
| ENSMUSG00000 | Gm11223 | -3.11E+00 | -4.12E+00 | -2.10E+00 | 2.846773 | -6.49E+00 | 4.95E-06 | 2.24E-03 | 4.213223 | -8.6477 | 1.24667891 | 0.03865355 | 3.22902915 | 1.524752578 | 0.92148254 | 4.74561156 | 4.757568734 | 4.11605873 | 3.237021006 | 4.65087791 |
| ENSMUSG00000 | Tinagl1 | -1.80E+00 | -2.39E+00 | -1.22E+00 | 5.176163 | -6.46E+00 | 5.31E-06 | 2.33E-03 | 4.17772 | -3.49361 | 3.122640253 | 5.12639621 | 4.419976927 | 3.734115645 | 5.30435805 | 6.05884921 | 6.070629897 | 5.98812953 | 6.424455776 | 5.51207737 |
| ENSMUSG00000 | Nkg7 | 1.12E+00 | 7.49E-01 | 1.49E+00 | 5.885028 | 6.38E+00 | 6.12E-06 | 2.45E-03 | 3.849591 | 2.170845 | 6.8449759 | 5.95154453 | 6.149100129 | 6.935795696 | 6.19709036 | 5.64871948 | 5.396343462 | 5.2030818 | 5.034998007 | 5.48862706 |
| ENSMUSG00000 | Kazald1 | -1.53E+00 | -2.03E+00 | -1.03E+00 | 4.441602 | -6.41E+00 | 5.85E-06 | 2.45E-03 | 4.139315 | -2.88592 | 2.879544694 | 3.84895056 | 3.578223385 | 3.746808059 | 4.10769831 | 5.32314361 | 5.095353941 | 5.03431382 | 6.044962394 | 4.75702465 |
| ENSMUSG00000 | Ncam1 | -3.16E+00 | -4.20E+00 | -2.11E+00 | 2.268986 | -6.38E+00 | 6.13E-06 | 2.45E-03 | 3.807573 | -8.91155 | 0.170593476 | -0.08072741 | 1.134875489 | 1.041093399 | 1.5371041 | 3.0832228 | 3.97758699 | 4.25462293 | 3.757608309 | 3.81388268 |
| ENSMUSG00000 | Mtss2 | -1.57E+00 | -2.10E+00 | -1.03E+00 | 3.459532 | -6.20E+00 | 8.85E-06 | 3.44E-03 | 3.781271 | -2.96335 | 2.831220247 | 3.17285553 | 2.825356181 | 1.826693768 | 3.01974311 | 3.94110122 | 4.271728776 | 4.25223831 | 4.202911251 | 4.25147272 |
| ENSMUSG00000 | Paqr7 | -7.80E-01 | -1.05E+00 | -5.08E-01 | 6.709672 | -6.05E+00 | 1.19E-05 | 4.49E-03 | 3.137455 | -1.71692 | 6.150689102 | 6.66676421 | 6.343694703 | 6.039804703 | 6.6578954 | 7.03885773 | 6.977054925 | 7.28074311 | 6.904263531 | 7.03694887 |
| ENSMUSG00000 | Serpinf1 | -8.96E-01 | -1.21E+00 | -5.78E-01 | 6.628291 | -5.94E+00 | 1.49E-05 | 5.21E-03 | 2.915685 | -1.86143 | 6.1478734 | 6.21210927 | 5.945995603 | 6.379755509 | 6.06269898 | 7.53601385 | 7.032047909 | 6.9313167 | 7.198454227 | 6.83664631 |
| ENSMUSG00000 | Maged2 | -6.82E-01 | -9.23E-01 | -4.40E-01 | 6.627621 | -5.95E+00 | 1.46E-05 | 5.21E-03 | 2.933645 | -1.60409 | 6.216668079 | 6.46536079 | 6.152724993 | 6.212925466 | 6.50382389 | 6.81937014 | 7.044290062 | 7.10853124 | 6.963501236 | 6.78900988 |
| ENSMUSG00000 | Siglecg | 1.24E+00 | 8.03E-01 | 1.68E+00 | 3.438496 | 5.94E+00 | 1.47E-05 | 5.21E-03 | 3.26809 | 2.368482 | 4.655029101 | 3.92866255 | 4.012508028 | 4.12288749 | 3.24173683 | 2.89947413 | 2.817205761 | 2.79071691 | 2.957735963 | 2.9590076 |
| ENSMUSG00000 | Ephb4 | -1.69E+00 | -2.30E+00 | -1.08E+00 | 2.625451 | -5.85E+00 | 1.75E-05 | 5.99E-03 | 3.097698 | -3.23186 | 1.869324852 | 1.54667036 | 1.486981733 | 1.675576035 | 2.6019124 | 3.25677889 | 3.618660082 | 3.58059585 | 3.506069542 | 3.11194116 |
| ENSMUSG00000 | Adam11 | -1.04E+00 | -1.41E+00 | -6.62E-01 | 4.128689 | -5.82E+00 | 1.88E-05 | 6.27E-03 | 2.996411 | -2.05196 | 3.233034051 | 4.03433566 | 3.686216433 | 3.655522808 | 3.52732308 | 4.70538032 | 4.813236859 | 4.57392772 | 4.457159941 | 4.60075249 |
| ENSMUSG00000 | Taldo1 | 5.41E-01 | 3.44E-01 | 7.38E-01 | 8.292598 | 5.79E+00 | 2.00E-05 | 6.52E-03 | 2.521893 | 1.454946 | 8.626748831 | 8.44747535 | 8.417490077 | 8.707106002 | 8.52736234 | 8.05477821 | 8.035968197 | 7.93494542 | 8.050269926 | 8.12383885 |
| ENSMUSG00000 | H19 | -3.33E+00 | -4.55E+00 | -2.11E+00 | 0.18722 | -5.76E+00 | 2.11E-05 | 6.72E-03 | 1.717889 | -10.0577 | -2.39234272 | -1.50296041 | -0.522236797 | -1.76626152 | -0.86499434 | 1.96228177 | 1.814031514 | 1.5748822 | 1.764516678 | 1.80527982 |
| ENSMUSG00000 | Ntf5 | -4.45E+00 | -6.09E+00 | -2.82E+00 | -1.53051 | -5.73E+00 | 2.25E-05 | 7.01E-03 | 1.08079 | -21.9052 | -5.85177434 | -2.9768916 | -2.348207397 | -4.08818962 | -3.67234926 | 0.66917651 | 0.781898893 | 0.12116423 | 1.853600221 | 0.20644515 |
| ENSMUSG00000 | Itsn1 | -9.17E-01 | -1.26E+00 | -5.78E-01 | 5.909803 | -5.70E+00 | 2.38E-05 | 7.25E-03 | 2.527645 | -1.88832 | 5.155953778 | 5.77390842 | 5.835766261 | 4.886224972 | 5.97474934 | 6.32723101 | 6.329835536 | 6.35332443 | 6.239320571 | 6.22171878 |
| ENSMUSG00000 | Vdr | -8.53E-01 | -1.17E+00 | -5.35E-01 | 5.602607 | -5.66E+00 | 2.62E-05 | 7.82E-03 | 2.455538 | -1.80596 | 4.943453629 | 5.51451616 | 5.683713057 | 5.074201711 | 4.63780906 | 6.08061765 | 6.000456675 | 6.22726246 | 5.925736838 | 5.93830524 |
| ENSMUSG00000 | Fyb | 7.47E-01 | 4.67E-01 | 1.03E+00 | 6.177173 | 5.62E+00 | 2.84E-05 | 8.17E-03 | 2.267458 | 1.678076 | 6.468371832 | 6.28531812 | 7.01534202 | 6.751287143 | 6.30138828 | 5.90543361 | 5.940755542 | 5.7469006 | 5.405753424 | 5.95118345 |
| ENSMUSG00000 | Hbb-bt | 1.33E+00 | 8.33E-01 | 1.83E+00 | 6.161815 | 5.61E+00 | 2.86E-05 | 8.17E-03 | 2.262897 | 2.520896 | 6.81366818 | 6.16571472 | 6.477948152 | 7.893995897 | 6.26779399 | 5.56455371 | 5.533403145 | 5.59961088 | 5.590950195 | 5.71051278 |
| ENSMUSG00000 | Smtnl1 | -1.70E+00 | -2.34E+00 | -1.06E+00 | 3.126239 | -5.59E+00 | 3.01E-05 | 8.43E-03 | 2.620835 | -3.24504 | 1.514547878 | 2.06750253 | 2.242209533 | 1.963472502 | 3.98728992 | 3.67905125 | 3.770481235 | 4.02526432 | 4.416802203 | 3.59577253 |
| ENSMUSG00000 | Rhbdf1 | -1.80E+00 | -2.48E+00 | -1.12E+00 | 3.322123 | -5.57E+00 | 3.15E-05 | 8.48E-03 | 2.578268 | -3.47418 | 1.287777016 | 3.2857253 | 3.250352707 | 1.858229342 | 2.45280587 | 3.94438008 | 4.188381631 | 4.2829351 | 4.754141679 | 3.91650287 |
| ENSMUSG00000 | Meg3 | -6.35E+00 | -8.75E+00 | -3.95E+00 | 2.343223 | -5.57E+00 | 3.11E-05 | 8.48E-03 | 1.941197 | -81.7308 | -1.60384682 | -3.8248885 | 1.885847941 | -0.3877499 | 0.29887238 | 5.993362 | 5.66310492 | 4.57006518 | 5.110592767 | 5.72686739 |
| ENSMUSG00000 | Tns1 | -7.88E-01 | -1.09E+00 | -4.86E-01 | 6.103418 | -5.51E+00 | 3.57E-05 | 9.44E-03 | 2.092036 | -1.72643 | 5.543224178 | 5.97504065 | 5.819438824 | 5.205665166 | 6.14367438 | 6.47738049 | 6.318119851 | 6.41645071 | 6.713367467 | 6.42181987 |
| ENSMUSG00000 | Pdgfd | -1.26E+00 | -1.74E+00 | -7.74E-01 | 3.486854 | -5.48E+00 | 3.77E-05 | 9.77E-03 | 2.37884 | -2.39058 | 2.52760403 | 3.24335236 | 2.795987329 | 3.463838993 | 1.85747168 | 4.09359662 | 4.222300457 | 4.16619181 | 4.137007471 | 4.36118989 |
| ENSMUSG00000 | Cracr2b | -1.34E+00 | -1.86E+00 | -8.19E-01 | 4.114932 | -5.44E+00 | 4.12E-05 | 1.05E-02 | 2.24868 | -2.52695 | 2.698510712 | 3.84311713 | 3.029168792 | 3.452261352 | 4.56926215 | 5.12116651 | 4.685006665 | 4.64701523 | 4.606528245 | 4.49728407 |
| ENSMUSG00000 | H2bc4 | 1.01E+00 | 6.13E-01 | 1.40E+00 | 5.557954 | 5.39E+00 | 4.53E-05 | 1.13E-02 | 1.890665 | 2.008001 | 6.30957173 | 5.58973755 | 5.513917086 | 5.94085829 | 6.6803872 | 4.66409435 | 4.812012235 | 5.05425786 | 6.005783641 | 5.00892311 |
| ENSMUSG00000 | Rtkn | -1.09E+00 | -1.51E+00 | -6.60E-01 | 4.938614 | -5.37E+00 | 4.71E-05 | 1.16E-02 | 1.992075 | -2.12279 | 4.084686935 | 5.04321793 | 4.474569094 | 3.898828001 | 4.6106094 | 5.26109787 | 5.461624968 | 5.69740593 | 5.69009285 | 5.16400988 |
| ENSMUSG00000 | Slco2a1 | 1.94E+00 | 1.18E+00 | 2.70E+00 | 2.263219 | 5.35E+00 | 4.89E-05 | 1.18E-02 | 2.163788 | 3.829704 | 2.810003761 | 3.64922301 | 2.916874837 | 3.340868351 | 2.77059423 | 1.49229874 | 0.177307585 | 1.30726053 | 1.836218143 | 2.33154037 |
| ENSMUSG00000 | Klk8 | -8.83E-01 | -1.23E+00 | -5.35E-01 | 5.033149 | -5.35E+00 | 4.96E-05 | 1.18E-02 | 1.913861 | -1.8439 | 4.626995283 | 3.97074859 | 4.647708755 | 4.631769551 | 4.9504347 | 5.42114866 | 5.37256888 | 5.24637927 | 5.939028949 | 5.52470958 |
| ENSMUSG00000 | Prss57 | 1.17E+00 | 7.03E-01 | 1.63E+00 | 4.073055 | 5.31E+00 | 5.35E-05 | 1.21E-02 | 1.924 | 2.2431 | 4.653640273 | 4.36639778 | 4.286280268 | 5.285965964 | 4.07852518 | 3.68983598 | 3.410598936 | 3.1780493 | 4.017715886 | 3.76354085 |
| ENSMUSG00000 | Cd69 | 8.91E-01 | 5.38E-01 | 1.25E+00 | 4.798792 | 5.31E+00 | 5.37E-05 | 1.21E-02 | 1.81977 | 1.854891 | 5.261316647 | 5.23310322 | 5.267171445 | 5.036793261 | 5.59396879 | 4.24486119 | 4.397934378 | 3.97119249 | 4.165124221 | 4.81645357 |
| ENSMUSG00000 | Gm10053 | 6.85E-01 | 4.13E-01 | 9.58E-01 | 6.815659 | 5.30E+00 | 5.45E-05 | 1.21E-02 | 1.560814 | 1.608224 | 7.566254813 | 6.70615759 | 7.124579427 | 7.333286173 | 6.81836125 | 6.52671462 | 6.531568441 | 6.46709331 | 6.650557263 | 6.43201516 |
| ENSMUSG00000 | Snrpn | -1.48E+00 | -2.06E+00 | -8.90E-01 | 2.860421 | -5.30E+00 | 5.45E-05 | 1.21E-02 | 2.063852 | -2.7829 | 1.632041441 | 2.09635739 | 1.857696901 | 2.343656169 | 3.188117 | 3.45347339 | 3.658139275 | 3.87217456 | 3.116922562 | 3.38563307 |
| ENSMUSG00000 | Pkn3 | -9.57E-01 | -1.34E+00 | -5.72E-01 | 4.59859 | -5.23E+00 | 6.26E-05 | 1.36E-02 | 1.741272 | -1.94085 | 4.23966105 | 4.497941 | 3.908322974 | 3.906163819 | 4.39481616 | 4.84525373 | 5.315596921 | 5.07805591 | 4.5718716 | 5.22822096 |
| ENSMUSG00000 | Tle2 | -8.44E-01 | -1.18E+00 | -5.04E-01 | 5.247591 | -5.23E+00 | 6.31E-05 | 1.36E-02 | 1.637808 | -1.79467 | 4.464507195 | 5.42919528 | 4.867435904 | 4.544805579 | 4.80095061 | 5.58817076 | 5.709863662 | 5.49897203 | 5.875569261 | 5.6964385 |
| ENSMUSG00000 | Gdf3 | 6.35E-01 | 3.78E-01 | 8.93E-01 | 5.983803 | 5.20E+00 | 6.73E-05 | 1.43E-02 | 1.422374 | 1.553443 | 6.31281838 | 6.22577644 | 6.141093224 | 6.536605838 | 6.02913049 | 5.53913466 | 5.647033023 | 5.59994851 | 5.890382542 | 5.91610339 |
| ENSMUSG00000 | Hid1 | -9.39E-01 | -1.32E+00 | -5.58E-01 | 5.381706 | -5.18E+00 | 6.96E-05 | 1.45E-02 | 1.526592 | -1.91742 | 4.43015569 | 5.64149391 | 4.619673828 | 4.613405643 | 5.49907776 | 5.66868099 | 5.905973619 | 5.79194875 | 5.844392075 | 5.80225666 |
| ENSMUSG00000 | Dlk1 | -3.77E+00 | -5.31E+00 | -2.23E+00 | -0.32963 | -5.17E+00 | 7.20E-05 | 1.46E-02 | 0.537626 | -13.6426 | -2.39234272 | -4.5618541 | -1.559711503 | -1.9727124 | -0.06354002 | 1.84086197 | 1.318399904 | 1.27002762 | 1.546711526 | 1.27782978 |
| ENSMUSG00000 | Gm42031 | -1.87E+00 | -2.63E+00 | -1.11E+00 | 2.261776 | -5.17E+00 | 7.20E-05 | 1.46E-02 | 1.781848 | -3.6533 | 2.41931513 | 0.58735107 | 1.463263633 | 1.3498814 | 0.41850417 | 3.02940184 | 3.332932629 | 2.95039611 | 3.439944283 | 3.62676546 |
| ENSMUSG00000 | Itgb2 | 6.95E-01 | 4.10E-01 | 9.79E-01 | 5.807476 | 5.15E+00 | 7.51E-05 | 1.48E-02 | 1.333659 | 1.618392 | 6.185429405 | 6.33170025 | 5.759841785 | 6.327200059 | 5.87493089 | 5.39799407 | 5.422632365 | 5.30783473 | 5.743108739 | 5.72408601 |
| ENSMUSG00000 | Cavin2 | -8.35E-01 | -1.18E+00 | -4.94E-01 | 4.699868 | -5.15E+00 | 7.48E-05 | 1.48E-02 | 1.546267 | -1.78391 | 4.420855448 | 4.13742915 | 3.805229481 | 4.364394835 | 4.90885132 | 5.09728355 | 5.119982756 | 5.03985466 | 4.826347899 | 5.2784515 |
| ENSMUSG00000 | Pvt1 | 1.53E+00 | 9.03E-01 | 2.16E+00 | 2.46722 | 5.14E+00 | 7.62E-05 | 1.48E-02 | 1.749434 | 2.886914 | 2.303043772 | 3.58280415 | 3.497784373 | 3.194126621 | 3.48060479 | 1.14665458 | 1.624107647 | 1.88915672 | 2.309099717 | 1.64481515 |
| ENSMUSG00000 | Ehd3 | -1.09E+00 | -1.54E+00 | -6.43E-01 | 4.823096 | -5.13E+00 | 7.83E-05 | 1.50E-02 | 1.511294 | -2.12978 | 3.516732125 | 4.45402552 | 4.497282653 | 3.954381766 | 5.27543376 | 4.95660983 | 5.303198684 | 5.44047638 | 5.603680338 | 5.22913459 |
| ENSMUSG00000 | Ncf4 | 7.21E-01 | 4.25E-01 | 1.02E+00 | 4.508061 | 5.12E+00 | 7.94E-05 | 1.50E-02 | 1.476031 | 1.648854 | 4.925480979 | 5.00736861 | 4.594307108 | 4.86697592 | 4.72425552 | 4.18790531 | 4.054917387 | 4.10407516 | 4.496419372 | 4.11890063 |
| ENSMUSG00000 | Plac8 | 1.21E+00 | 7.10E-01 | 1.71E+00 | 6.286432 | 5.11E+00 | 8.20E-05 | 1.52E-02 | 1.191163 | 2.309946 | 6.716419317 | 6.45796826 | 6.83057061 | 7.531571845 | 6.33691952 | 5.97476225 | 5.180392245 | 5.54353376 | 5.898779481 | 6.39340266 |
| ENSMUSG00000 | P2ry14 | 6.10E-01 | 3.58E-01 | 8.61E-01 | 5.837574 | 5.11E+00 | 8.23E-05 | 1.52E-02 | 1.241206 | 1.526067 | 6.234693657 | 5.80191564 | 6.19833634 | 6.141029765 | 6.40473348 | 5.45312815 | 5.579472794 | 5.61487468 | 5.601109834 | 5.34644129 |
| ENSMUSG00000 | Ppa1 | 5.43E-01 | 3.18E-01 | 7.67E-01 | 7.648284 | 5.08E+00 | 8.65E-05 | 1.55E-02 | 1.054965 | 1.45654 | 8.052201628 | 7.91261237 | 7.808565194 | 7.931285282 | 7.78554457 | 7.57249558 | 7.344347608 | 7.40777222 | 7.536153334 | 7.13185983 |
| ENSMUSG00000 | Syde1 | -8.47E-01 | -1.20E+00 | -4.96E-01 | 4.85594 | -5.08E+00 | 8.65E-05 | 1.55E-02 | 1.373439 | -1.79829 | 4.044558067 | 4.9649581 | 4.384801759 | 4.510483263 | 4.34668541 | 5.52714657 | 5.304307588 | 5.18682895 | 4.952703886 | 5.33692245 |
| ENSMUSG00000 | Nipal2 | -2.79E+00 | -3.95E+00 | -1.63E+00 | 0.826134 | -5.07E+00 | 8.82E-05 | 1.56E-02 | 1.166254 | -6.91653 | -1.09688683 | -1.01753358 | 0.792273826 | -1.02929593 | 0.02809046 | 2.46058988 | 2.182239881 | 2.42626771 | 2.016790389 | 1.49880013 |
| ENSMUSG00000 | Pde1b | -1.02E+00 | -1.44E+00 | -5.94E-01 | 4.800099 | -5.06E+00 | 9.03E-05 | 1.58E-02 | 1.361265 | -2.02344 | 3.638073624 | 5.0374389 | 3.877109604 | 4.291188749 | 4.52240759 | 5.27055957 | 5.204834932 | 5.38604874 | 5.677520397 | 5.09580582 |
| ENSMUSG00000 | Lat | -5.77E-01 | -8.18E-01 | -3.37E-01 | 6.725237 | -5.05E+00 | 9.16E-05 | 1.58E-02 | 1.065429 | -1.49199 | 6.168858425 | 6.63105512 | 6.473421656 | 6.255736321 | 6.64800002 | 6.90642587 | 6.880215559 | 7.03344614 | 7.304564487 | 6.95064525 |
| ENSMUSG00000 | Xlr4a | 1.10E+00 | 6.43E-01 | 1.56E+00 | 3.815828 | 5.05E+00 | 9.25E-05 | 1.58E-02 | 1.434387 | 2.147676 | 4.446311149 | 3.85197126 | 3.835351297 | 4.510929772 | 5.20743001 | 3.35056399 | 2.984831894 | 3.04936288 | 3.420597619 | 3.50093271 |
| ENSMUSG00000 | Krt80 | 1.31E+00 | 7.62E-01 | 1.86E+00 | 3.860251 | 5.04E+00 | 9.52E-05 | 1.61E-02 | 1.389551 | 2.47821 | 4.483839328 | 4.27883992 | 3.419639978 | 5.223664634 | 4.93081713 | 3.25493148 | 3.211388196 | 3.11151406 | 3.503869472 | 3.18400382 |
| ENSMUSG00000 | Runx1t1 | -1.74E+00 | -2.47E+00 | -1.01E+00 | 4.034415 | -5.00E+00 | 1.02E-04 | 1.70E-02 | 1.407133 | -3.33815 | 2.208921595 | 4.02185552 | 3.955573351 | 1.842547719 | 4.32426545 | 4.68592358 | 4.525671958 | 4.99777015 | 5.172934318 | 4.60868486 |
| ENSMUSG00000 | Vipr2 | -6.07E-01 | -8.64E-01 | -3.51E-01 | 5.517937 | -4.98E+00 | 1.07E-04 | 1.77E-02 | 1.0469 | -1.52361 | 5.349737008 | 5.20025106 | 4.939714813 | 5.255366256 | 5.18500975 | 5.8956681 | 5.695273522 | 5.78866149 | 5.994927274 | 5.87476492 |
| ENSMUSG00000 | Smim24 | -8.41E-01 | -1.20E+00 | -4.84E-01 | 5.314847 | -4.95E+00 | 1.13E-04 | 1.84E-02 | 1.043663 | -1.79143 | 4.356460022 | 5.08740208 | 5.281811465 | 4.797506755 | 4.7730795 | 5.86398747 | 5.514471232 | 5.76543854 | 6.188856841 | 5.5194596 |
| ENSMUSG00000 | Cox5a | 5.63E-01 | 3.23E-01 | 8.03E-01 | 7.398714 | 4.94E+00 | 1.16E-04 | 1.87E-02 | 0.768299 | 1.477134 | 7.817828272 | 7.32769043 | 7.71592539 | 7.868580982 | 7.39435886 | 7.1889458 | 7.167109146 | 7.00842459 | 7.428937918 | 7.06933491 |
| ENSMUSG00000 | Gm10131 | 5.86E-01 | 3.36E-01 | 8.37E-01 | 7.117535 | 4.93E+00 | 1.20E-04 | 1.89E-02 | 0.748243 | 1.501187 | 7.578474342 | 7.08443759 | 7.677858253 | 7.538051767 | 7.00388288 | 7.02823963 | 6.748959463 | 6.85861195 | 6.822231838 | 6.83460589 |
| ENSMUSG00000 | Ccnd1 | 6.22E-01 | 3.56E-01 | 8.88E-01 | 5.680145 | 4.93E+00 | 1.20E-04 | 1.89E-02 | 0.872942 | 1.538771 | 6.225782455 | 6.0697959 | 5.989138791 | 6.040131281 | 5.44861432 | 5.53869002 | 5.510858753 | 5.26969755 | 5.429955799 | 5.27878711 |
| ENSMUSG00000 | Hdac7 | -5.91E-01 | -8.44E-01 | -3.37E-01 | 5.66277 | -4.91E+00 | 1.24E-04 | 1.90E-02 | 0.87907 | -1.50582 | 5.321278121 | 5.30696846 | 5.238802932 | 5.374653435 | 5.61628615 | 5.8038849 | 5.978650532 | 6.05669231 | 6.119601194 | 5.81087751 |
| ENSMUSG00000 | Tmem47 | -3.15E+00 | -4.50E+00 | -1.80E+00 | -0.03145 | -4.91E+00 | 1.24E-04 | 1.90E-02 | 0.636639 | -8.86328 | -1.94488374 | -0.6549635 | -1.559711503 | -0.3877499 | -4.40931486 | 1.3212532 | 1.970476601 | 1.7473494 | 1.412410434 | 2.19067783 |
| ENSMUSG00000 | Lgals1 | 9.73E-01 | 5.56E-01 | 1.39E+00 | 4.972068 | 4.91E+00 | 1.25E-04 | 1.90E-02 | 0.955335 | 1.963263 | 5.04076848 | 4.82974753 | 5.510338806 | 5.754684485 | 5.87106929 | 4.62388414 | 4.474318048 | 4.11985685 | 5.290872622 | 4.20513776 |
| ENSMUSG00000 | Rexo2 | 5.20E-01 | 2.94E-01 | 7.45E-01 | 6.587546 | 4.86E+00 | 1.39E-04 | 2.06E-02 | 0.636256 | 1.433673 | 7.001340247 | 6.66916703 | 6.767663859 | 6.984836157 | 6.6605824 | 6.33224582 | 6.433790957 | 6.19472371 | 6.450392565 | 6.38071955 |
| ENSMUSG00000 | Mturn | -1.16E+00 | -1.67E+00 | -6.56E-01 | 3.489556 | -4.84E+00 | 1.44E-04 | 2.06E-02 | 1.097691 | -2.23605 | 2.492521571 | 3.68131989 | 3.170156064 | 2.33247243 | 3.06371436 | 3.8841894 | 4.13086624 | 4.0529295 | 4.090115755 | 3.99727761 |
| ENSMUSG00000 | Cux2 | -3.04E+00 | -4.36E+00 | -1.73E+00 | 0.514915 | -4.86E+00 | 1.38E-04 | 2.06E-02 | 0.639323 | -8.25306 | -0.89757803 | -0.16953667 | -1.415321593 | -1.9727124 | -0.16138734 | 2.65723689 | 1.801264218 | 1.48024533 | 1.746020334 | 2.08091426 |
| ENSMUSG00000 | Glipr1 | 4.65E-01 | 2.63E-01 | 6.67E-01 | 6.712646 | 4.84E+00 | 1.43E-04 | 2.06E-02 | 0.592746 | 1.380434 | 6.926590317 | 6.93998309 | 7.004650619 | 7.05413021 | 6.8470747 | 6.43267564 | 6.533871104 | 6.57345502 | 6.264766627 | 6.54926576 |
| ENSMUSG00000 | Il11ra1 | -5.49E-01 | -7.88E-01 | -3.10E-01 | 7.4576 | -4.85E+00 | 1.43E-04 | 2.06E-02 | 0.567673 | -1.46304 | 7.133356037 | 7.41982259 | 6.97869188 | 7.01248586 | 7.43114838 | 7.73086014 | 7.779526976 | 7.69476752 | 7.929406955 | 7.46593053 |
| ENSMUSG00000 | Airn | -3.05E+00 | -4.38E+00 | -1.73E+00 | -0.3364 | -4.84E+00 | 1.43E-04 | 2.06E-02 | 0.380551 | -8.3012 | -1.32821238 | -2.239926 | -0.678355999 | -1.42522461 | -4.40931486 | 1.49229874 | 1.471298804 | 1.39634407 | 1.388759166 | 0.96828541 |
| ENSMUSG00000 | Slc25a5 | 6.13E-01 | 3.46E-01 | 8.80E-01 | 10.48378 | 4.83E+00 | 1.47E-04 | 2.07E-02 | 0.466501 | 1.529476 | 11.15351862 | 10.589547 | 10.38775396 | 10.97134779 | 10.4775055 | 10.117969 | 10.19356218 | 10.0883604 | 10.61999701 | 10.238264 |
| ENSMUSG00000 | Alcam | -9.77E-01 | -1.40E+00 | -5.51E-01 | 4.316817 | -4.83E+00 | 1.47E-04 | 2.07E-02 | 0.945465 | -1.96899 | 3.467897784 | 3.63618161 | 4.168208952 | 3.856278435 | 4.34890836 | 4.92873872 | 4.579005373 | 4.77834026 | 4.256165557 | 5.14844959 |
| ENSMUSG00000 | Gng11 | -6.54E-01 | -9.40E-01 | -3.68E-01 | 5.524755 | -4.81E+00 | 1.53E-04 | 2.11E-02 | 0.689033 | -1.57328 | 4.871886608 | 5.29768069 | 5.276504994 | 5.256845944 | 5.46975779 | 6.08878948 | 5.872798087 | 5.77460647 | 5.603680338 | 5.73499835 |
| ENSMUSG00000 | Rian | -3.76E+00 | -5.40E+00 | -2.11E+00 | -0.12431 | -4.81E+00 | 1.53E-04 | 2.11E-02 | 0.419414 | -13.5022 | -1.94488374 | -2.68738498 | 0.300885441 | -0.5438691 | -5.99427736 | 2.19425412 | 2.627727963 | 1.76083263 | 1.00905474 | 2.03455863 |
| ENSMUSG00000 | Rras | -1.23E+00 | -1.77E+00 | -6.92E-01 | 3.208327 | -4.80E+00 | 1.57E-04 | 2.14E-02 | 1.045166 | -2.3512 | 1.695120123 | 3.03060294 | 2.396932128 | 2.588942727 | 3.30721884 | 3.62286131 | 3.843908555 | 3.66906464 | 4.199516673 | 3.72909774 |

| | | | | | | | | | | | | | | | | | | | | |
|---|---|---|---|---|---|---|---|---|---|---|---|---|---|---|---|---|---|---|---|---|
| ENSMUSG00000( | Abcg3 | -7.69E-01 | -1.11E+00 | -4.30E-01 | 5.623326 | -4.78E+00 | 1.63E-04 | 2.20E-02 | 0.60968 | -1.70444 | 5.04228551 | 5.87520437 | 5.451515753 | 5.197983065 | 4.51054163 | 6.27284361 | 5.985720848 | 5.98741284 | 5.833483176 | 6.07627365 |
| ENSMUSG00000( | Mpped2 | -2.59E+00 | -3.74E+00 | -1.44E+00 | 1.126428 | -4.76E+00 | 1.71E-04 | 2.26E-02 | 0.796546 | -6.02531 | -0.89757803 | 1.2368877 | 1.203588239 | -0.628758 | -2.08738676 | 2.73502905 | 2.810850266 | 2.30265862 | 2.614554172 | 1.97443764 |
| ENSMUSG00000( | Xlr4b | 9.91E-01 | 5.52E-01 | 1.43E+00 | 3.985167 | 4.76E+00 | 1.72E-04 | 2.26E-02 | 0.795587 | 1.986991 | 4.637792682 | 4.21051547 | 4.001871896 | 4.521973979 | 5.06566191 | 3.14636988 | 3.856953491 | 3.33740539 | 3.895825134 | 3.1772988 |
| ENSMUSG00000( | Rbm47 | 9.20E-01 | 5.13E-01 | 1.33E+00 | 3.906917 | 4.76E+00 | 1.73E-04 | 2.26E-02 | 0.78981 | 1.892302 | 4.265868765 | 3.98889269 | 4.531097366 | 4.638596196 | 4.67726371 | 3.65122983 | 3.316157954 | 3.37238934 | 2.759914776 | 3.86775853 |
| ENSMUSG00000( | Obsl1 | -9.92E-01 | -1.43E+00 | -5.52E-01 | 4.007275 | -4.75E+00 | 1.75E-04 | 2.27E-02 | 0.834609 | -1.98899 | 3.099510378 | 3.47706489 | 3.775218019 | 3.278132596 | 3.90809776 | 4.13436165 | 4.577134173 | 4.43154651 | 5.092862874 | 4.29881929 |
| ENSMUSG00000( | Lancl3 | -2.01E+00 | -2.90E+00 | -1.12E+00 | 1.620815 | -4.74E+00 | 1.77E-04 | 2.28E-02 | 0.924867 | -4.02137 | 1.067088901 | 1.43061223 | 1.501428004 | 0.701887313 | -2.29383764 | 2.8927009 | 2.725575286 | 2.82885546 | 2.334082074 | 3.01976163 |
| ENSMUSG00000( | Cd68 | 8.34E-01 | 4.61E-01 | 1.21E+00 | 4.393386 | 4.70E+00 | 1.94E-04 | 2.42E-02 | 0.620225 | 1.782754 | 4.612771414 | 4.87901507 | 4.753648929 | 4.624910449 | 4.86448074 | 4.29965367 | 3.723884958 | 3.65107569 | 4.70710589 | 3.81730951 |
| ENSMUSG00000( | Plekhg1 | -1.57E+00 | -2.27E+00 | -8.66E-01 | 1.781654 | -4.71E+00 | 1.91E-04 | 2.42E-02 | 0.868608 | -2.9608 | 0.591169159 | 1.44564044 | 0.668094415 | 1.068314868 | 1.06100508 | 2.6652076 | 2.591109371 | 2.37458355 | 2.900348481 | 2.45106751 |
| ENSMUSG00000: | Gm29946 | 6.49E-01 | 3.59E-01 | 9.40E-01 | 5.12424 | 4.71E+00 | 1.92E-04 | 2.42E-02 | 0.491534 | 1.568492 | 5.069322751 | 5.54449073 | 5.842066249 | 5.409662219 | 5.48398667 | 4.82405024 | 4.844686498 | 4.71740802 | 4.690274257 | 4.81645357 |
| ENSMUSG00000( | Scrn1 | -1.16E+00 | -1.68E+00 | -6.41E-01 | 3.563186 | -4.70E+00 | 1.95E-04 | 2.42E-02 | 0.793922 | -2.23856 | 2.292883906 | 3.67495739 | 2.781075619 | 2.885268594 | 3.11685831 | 4.075767 | 3.81879027 | 4.15602299 | 4.504694892 | 4.32553663 |
| ENSMUSG00000( | 1700003E16Rik | -3.74E+00 | -5.42E+00 | -2.06E+00 | 0.385796 | -4.69E+00 | 2.01E-04 | 2.47E-02 | 0.382229 | -13.377 | -3.52984624 | 0.56742892 | -3.000284094 | 1.338075137 | -4.40931486 | 2.77976737 | 2.201802147 | 2.86126272 | 2.624714038 | 2.42435018 |
| ENSMUSG00000: | Gm37795 | 1.62E+00 | 8.90E-01 | 2.35E+00 | 1.989324 | 4.68E+00 | 2.02E-04 | 2.47E-02 | 0.84664 | 3.069421 | 3.052107509 | 2.79862724 | 2.680201018 | 2.251660385 | 3.42988893 | 0.60432536 | 1.518864487 | 1.60509681 | 1.746020334 | 0.20644515 |
| ENSMUSG00000( | Cfp | 9.97E-01 | 5.47E-01 | 1.45E+00 | 4.98064 | 4.67E+00 | 2.09E-04 | 2.53E-02 | 0.425281 | 1.996407 | 5.610216614 | 5.41685636 | 4.546610366 | 6.015535304 | 5.38672475 | 4.83292293 | 4.308283575 | 4.27120581 | 4.815301176 | 4.60273968 |
| ENSMUSG00000( | A530040E14Rik | 1.36E+00 | 7.45E-01 | 1.98E+00 | 3.028498 | 4.65E+00 | 2.15E-04 | 2.58E-02 | 0.704202 | 2.568714 | 3.175799968 | 3.94858043 | 3.182691903 | 4.470741242 | 3.2788655 | 2.7306299 | 1.837607341 | 2.53274768 | 2.55204497 | 2.57527422 |
| ENSMUSG00000( | Enho | -3.52E+00 | -5.11E+00 | -1.92E+00 | -0.98473 | -4.64E+00 | 2.22E-04 | 2.63E-02 | -0.2306 | -11.4397 | -1.45945691 | -2.44637688 | -5.807639016 | -2.50322712 | -1.35042117 | 0.70054068 | 0.857847746 | 0.61255262 | 0.73668247 | 0.81216621 |
| ENSMUSG00000( | Ephb6 | -1.12E+00 | -1.62E+00 | -6.09E-01 | 3.493103 | -4.64E+00 | 2.23E-04 | 2.63E-02 | 0.671811 | -2.16732 | 2.706646377 | 3.33498284 | 2.704113638 | 2.748912647 | 3.04189625 | 3.67115687 | 3.806065204 | 4.21353838 | 4.437122188 | 4.26659281 |
| ENSMUSG00000( | Fgfr3 | -1.50E+00 | -2.18E+00 | -8.17E-01 | 3.221567 | -4.63E+00 | 2.29E-04 | 2.67E-02 | 0.688384 | -2.82655 | 2.68350104 | 2.92196168 | 2.688216011 | 2.006327981 | 1.81950383 | 4.0332871 | 3.448962474 | 4.49340979 | 4.229784112 | 3.89071098 |
| ENSMUSG00000( | Hnf4a | -9.70E-01 | -1.42E+00 | -5.24E-01 | 4.076401 | -4.58E+00 | 2.52E-04 | 2.87E-02 | 0.434876 | -1.9589 | 3.259361334 | 3.56226722 | 3.666066733 | 4.279589128 | 3.06911772 | 4.68983598 | 4.764738342 | 4.79153825 | 4.36319746 | 4.31829902 |
| ENSMUSG00000( | Plxdc2 | -7.66E-01 | -1.12E+00 | -4.14E-01 | 6.153148 | -4.59E+00 | 2.49E-04 | 2.87E-02 | 0.122341 | -1.70081 | 5.668353214 | 6.41852353 | 6.095112929 | 5.267161478 | 5.70399322 | 6.61747955 | 6.582040906 | 6.64085746 | 6.17941059 | 6.35854625 |
| ENSMUSG00000( | Pafah2 | 5.52E-01 | 2.98E-01 | 8.05E-01 | 7.086236 | 4.58E+00 | 2.52E-04 | 2.87E-02 | 0.00552 | 1.465809 | 7.092022483 | 7.57417575 | 7.534296711 | 7.148023937 | 7.67737427 | 6.84152158 | 6.61615639 | 6.91746253 | 6.661880442 | 6.79944376 |
| ENSMUSG00000( | Cav2 | -1.37E+00 | -2.00E+00 | -7.36E-01 | 3.431511 | -4.56E+00 | 2.64E-04 | 2.89E-02 | 0.500823 | -2.57945 | 3.0700666 | 3.18185833 | 2.337019227 | 3.206431131 | 1.83227113 | 4.60121275 | 4.635840608 | 3.99420706 | 3.642323438 | 3.81388268 |
| ENSMUSG00000( | Gfi1 | 2.12E+00 | 1.14E+00 | 3.10E+00 | 1.618052 | 4.56E+00 | 2.62E-04 | 2.89E-02 | 0.603372 | 4.343823 | 2.073038167 | 2.04794026 | 2.655885357 | 2.616866728 | 4.199248 | 1.70827633 | -0.081779636 | 0.7808241 | 1.128107412 | -0.947883 |
| ENSMUSG00000( | Dock10 | 6.19E-01 | 3.34E-01 | 9.04E-01 | 5.134899 | 4.57E+00 | 2.56E-04 | 2.89E-02 | 0.194388 | 1.535507 | 5.608169512 | 5.57385519 | 5.696683432 | 5.377376787 | 5.08587395 | 4.78800352 | 4.843130192 | 4.98633714 | 4.360160206 | 5.02939692 |
| ENSMUSG00000( | H2-Eb1 | 1.07E+00 | 5.75E-01 | 1.56E+00 | 5.437662 | 4.56E+00 | 2.62E-04 | 2.89E-02 | 0.140551 | 2.096125 | 5.403844413 | 6.37796996 | 5.850126019 | 6.007646916 | 5.80864284 | 5.68347294 | 4.60861914 | 4.51148898 | 5.622814505 | 4.50199783 |
| ENSMUSG00000( | Gm20427 | -1.63E+00 | -2.39E+00 | -8.79E-01 | 3.252033 | -4.56E+00 | 2.62E-04 | 2.89E-02 | 0.553015 | -3.10252 | 1.639918451 | 2.90303195 | 1.903718222 | 3.117294129 | 2.32729002 | 3.42253654 | 4.530379467 | 4.20419256 | 4.62860659 | 3.84335973 |
| ENSMUSG00000( | Rab19 | 1.29E+00 | 6.95E-01 | 1.89E+00 | 2.756101 | 4.55E+00 | 2.66E-04 | 2.89E-02 | 0.54959 | 2.450716 | 3.184399276 | 3.42493605 | 3.710030372 | 2.877594667 | 3.49155195 | 2.25413901 | 1.388362463 | 2.40065706 | 3.066085685 | 1.76325661 |
| ENSMUSG00000( | Rorc | -3.26E+00 | -4.77E+00 | -1.75E+00 | 1.574238 | -4.55E+00 | 2.71E-04 | 2.92E-02 | 0.400976 | -9.58374 | -2.68184934 | 0.74800117 | -0.598185651 | 0.054768336 | 3.27951824 | 3.58974204 | 2.932926863 | 3.29108924 | 2.989532182 | 2.13683968 |
| ENSMUSG00000( | Timm23 | 4.75E-01 | 2.54E-01 | 6.96E-01 | 6.294308 | 4.53E+00 | 2.78E-04 | 2.94E-02 | -0.03362 | 1.390027 | 6.620540805 | 6.34321194 | 6.544062184 | 6.504248731 | 6.50474133 | 5.98060297 | 5.934623153 | 6.1051723 | 6.345668314 | 6.06020942 |
| ENSMUSG00000( | Higd1a | 5.73E-01 | 3.07E-01 | 8.39E-01 | 6.007214 | 4.53E+00 | 2.78E-04 | 2.94E-02 | -0.00562 | 1.487394 | 6.505612716 | 5.90032095 | 6.185072775 | 6.470256607 | 6.17067355 | 5.61942956 | 5.786181623 | 5.62113487 | 6.04877805 | 5.76467728 |
| ENSMUSG00000( | Clca3a1 | -1.11E+00 | -1.63E+00 | -5.95E-01 | 3.29895 | -4.53E+00 | 2.79E-04 | 2.94E-02 | 0.479783 | -2.16143 | 2.510169437 | 3.117626 | 1.848285197 | 2.661791512 | 3.82430482 | 3.8097754 | 3.757122548 | 3.76585653 | 3.820835111 | 3.87372847 |
| ENSMUSG00000( | Tnfaip8l2 | 8.23E-01 | 4.40E-01 | 1.21E+00 | 4.066048 | 4.53E+00 | 2.82E-04 | 2.95E-02 | 0.290668 | 1.768868 | 4.047582586 | 4.52118194 | 4.350970672 | 4.758345485 | 4.60098012 | 3.43517747 | 3.62591894 | 3.51845133 | 3.78956805 | 4.01230582 |
| ENSMUSG00000( | Cldn5 | -4.41E+00 | -6.46E+00 | -2.35E+00 | -0.74614 | -4.52E+00 | 2.87E-04 | 2.98E-02 | -0.35572 | -21.2463 | -1.7643115 | -6.1468166 | -1.900748421 | -3.35122402 | -2.29383764 | 2.33630755 | 0.906346264 | 0.55179859 | 3.005171306 | 1.19595277 |
| ENSMUSG00000( | Aplp2 | -4.91E-01 | -7.21E-01 | -2.62E-01 | 7.556527 | -4.51E+00 | 2.94E-04 | 3.03E-02 | -0.16679 | -1.40589 | 7.208414942 | 7.69563197 | 7.396473572 | 7.212353611 | 6.95354219 | 7.8595439 | 7.754736413 | 7.86760553 | 7.897339731 | 7.71962919 |
| ENSMUSG00000( | Adamtsl4 | -2.45E+00 | -3.61E+00 | -1.30E+00 | 0.345664 | -4.49E+00 | 3.09E-04 | 3.11E-02 | 0.136722 | -5.47737 | -1.09688683 | -0.72055184 | -0.450087012 | -0.46369875 | -2.29383764 | 1.64519764 | 1.127034379 | 1.98418333 | 1.800812807 | 1.92446918 |
| ENSMUSG00000( | Trim47 | -7.33E-01 | -1.08E+00 | -3.89E-01 | 4.984557 | -4.49E+00 | 3.09E-04 | 3.11E-02 | 0.092865 | -1.66252 | 3.985853597 | 4.54055909 | 4.840718566 | 4.65888443 | 5.15911 | 5.2913088 | 5.188994371 | 5.2764958 | 5.472007705 | 5.43163512 |
| ENSMUSG00000( | Eng | -5.05E-01 | -7.43E-01 | -2.68E-01 | 8.033772 | -4.48E+00 | 3.12E-04 | 3.11E-02 | -0.24534 | -1.41961 | 7.490439259 | 8.10311163 | 7.608498752 | 7.835756811 | 7.84954557 | 8.2527346 | 8.389495792 | 8.24408599 | 8.449410441 | 8.11464194 |
| ENSMUSG00000( | Pnpla8 | -6.14E-01 | -9.02E-01 | -3.25E-01 | 5.597886 | -4.48E+00 | 3.12E-04 | 3.11E-02 | -0.0397 | -1.53009 | 5.329999102 | 5.25991989 | 5.332551687 | 5.293353333 | 5.61351528 | 6.12824777 | 6.025373111 | 5.89490071 | 5.256982254 | 5.84401507 |
| ENSMUSG00000( | C130026I21Rik | 1.55E+00 | 8.24E-01 | 2.29E+00 | 2.175795 | 4.48E+00 | 3.13E-04 | 3.11E-02 | 0.427904 | 2.937541 | 2.356606357 | 3.20553587 | 3.446302991 | 3.225177179 | 2.75411678 | 1.88698671 | 2.092658219 | 0.47239924 | 0.770019654 | 1.54814219 |
| ENSMUSG00000( | Slc22a17 | -7.73E-01 | -1.14E+00 | -4.08E-01 | 4.433897 | -4.46E+00 | 3.25E-04 | 3.13E-02 | 0.132776 | -1.70852 | 3.481381014 | 4.31568568 | 4.082625261 | 4.217112158 | 4.14143193 | 4.94846819 | 4.853989239 | 4.63377589 | 4.810858693 | 4.85364284 |
| ENSMUSG00000( | Mecr | 5.24E-01 | 2.76E-01 | 7.71E-01 | 5.407648 | 4.46E+00 | 3.26E-04 | 3.13E-02 | -0.07745 | 1.437451 | 5.689806323 | 5.4655912 | 5.659457018 | 5.692623516 | 5.69441007 | 5.11630516 | 5.091426456 | 5.00487004 | 5.42772349 | 5.23427103 |
| ENSMUSG00000( | Snrpa1 | 4.42E-01 | 2.34E-01 | 6.50E-01 | 7.441164 | 4.47E+00 | 3.18E-04 | 3.13E-02 | -0.24831 | 1.358722 | 7.842039281 | 7.52638204 | 7.697552264 | 7.692760181 | 7.64811285 | 7.13669248 | 7.354488173 | 7.26633956 | 6.97645146 | 7.27081694 |
| ENSMUSG00000( | Chchd4 | 4.90E-01 | 2.59E-01 | 7.22E-01 | 6.153445 | 4.46E+00 | 3.25E-04 | 3.13E-02 | -0.1808 | 1.404687 | 6.620154851 | 6.38041603 | 6.239825757 | 6.428495331 | 6.40473348 | 5.99782678 | 5.973680614 | 6.01580628 | 5.716042704 | 5.75747069 |
| ENSMUSG00000( | Cyp26b1 | -2.60E+00 | -3.83E+00 | -1.38E+00 | 2.558265 | -4.47E+00 | 3.21E-04 | 3.13E-02 | 0.422647 | -6.07088 | -1.60384682 | 2.06750253 | 1.857696901 | 1.842547719 | 2.16054075 | 4.45255257 | 3.615016911 | 3.17503111 | 4.111947622 | 3.90366456 |
| ENSMUSG00000( | Nhp2 | 4.61E-01 | 2.43E-01 | 6.79E-01 | 7.117252 | 4.45E+00 | 3.34E-04 | 3.13E-02 | -0.28008 | 1.376502 | 7.540945568 | 7.20260287 | 7.235900983 | 7.327728164 | 7.20186367 | 6.80591416 | 6.867257704 | 6.8096933 | 7.29704012 | 6.88357749 |
| ENSMUSG00000( | Cfl1 | 4.09E-01 | 2.16E-01 | 6.03E-01 | 10.28743 | 4.46E+00 | 3.29E-04 | 3.13E-02 | -0.3401 | 1.327912 | 10.65059972 | 10.3567657 | 10.35845461 | 10.54313184 | 10.4150781 | 10.0794289 | 10.10226485 | 9.97622905 | 10.26192259 | 10.130449 |
| ENSMUSG00000( | Osm | 9.39E-01 | 4.94E-01 | 1.38E+00 | 3.576109 | 4.45E+00 | 3.36E-04 | 3.13E-02 | 0.184403 | 1.916773 | 4.174749106 | 4.29301429 | 4.00774428 | 4.10984609 | 4.16937232 | 3.15856135 | 3.109016544 | 3.2196541 | 1.782778884 | 3.7363566 |
| ENSMUSG00000( | Mfap2 | -1.29E+00 | -1.90E+00 | -6.79E-01 | 3.565666 | -4.45E+00 | 3.36E-04 | 3.13E-02 | 0.272171 | -2.44461 | 1.5491051 | 3.58619873 | 2.846997012 | 3.368507033 | 3.00290212 | 4.06375698 | 4.285545594 | 4.15602299 | 4.806402489 | 3.99122222 |
| ENSMUSG00000( | Igf2r | -8.45E-01 | -1.25E+00 | -4.44E-01 | 4.200407 | -4.44E+00 | 3.40E-04 | 3.14E-02 | 0.129969 | -1.79585 | 3.693190096 | 3.47706489 | 3.942230411 | 3.954381766 | 3.73873796 | 4.26072968 | 4.659024093 | 4.59666272 | 4.737853013 | 4.9441946 |
| ENSMUSG00000( | Leng8 | -5.67E-01 | -8.35E-01 | -2.98E-01 | 6.994698 | -4.44E+00 | 3.38E-04 | 3.14E-02 | -0.27363 | -1.48108 | 6.611591658 | 7.27202163 | 6.695714689 | 6.40711247 | 6.73048374 | 7.22810681 | 7.291386467 | 7.22387105 | 7.125851438 | 7.36084355 |
| ENSMUSG00000( | Psmc5 | 4.54E-01 | 2.38E-01 | 6.70E-01 | 7.321178 | 4.43E+00 | 3.47E-04 | 3.17E-02 | -0.32797 | 1.369777 | 7.719622803 | 7.40019841 | 7.347653586 | 7.549189984 | 7.53478711 | 7.11102622 | 7.019011936 | 6.99509866 | 7.445922969 | 7.08926538 |
| ENSMUSG00000( | Srgap3 | -8.53E-01 | -1.26E+00 | -4.47E-01 | 4.481108 | -4.42E+00 | 3.53E-04 | 3.19E-02 | 0.055955 | -1.80586 | 4.068578519 | 4.0718435 | 4.322931547 | 3.677787063 | 4.18935802 | 4.79346828 | 5.141668236 | 4.53129293 | 5.130783199 | 4.88337316 |
| ENSMUSG00000( | Psme2 | 4.42E-01 | 2.32E-01 | 6.52E-01 | 7.436634 | 4.43E+00 | 3.51E-04 | 3.19E-02 | -0.34806 | 1.358278 | 7.91713777 | 7.56113132 | 7.534032686 | 7.726977497 | 7.46915761 | 7.3288596 | 7.267575199 | 7.21695109 | 7.202351695 | 7.1421611 |
| ENSMUSG00000( | Evc2 | -1.42E+00 | -2.09E+00 | -7.40E-01 | 2.651739 | -4.41E+00 | 3.63E-04 | 3.26E-02 | 0.288322 | -2.66817 | 2.465638277 | 2.77502434 | 2.006142175 | 1.810663658 | 0.1142471 | 3.23016014 | 3.563022273 | 3.71134444 | 3.637314075 | 3.2038292 |
| ENSMUSG00000( | Exoc6b | -7.45E-01 | -1.10E+00 | -3.88E-01 | 5.35549 | -4.39E+00 | 3.77E-04 | 3.36E-02 | -0.16843 | -1.67546 | 4.626995283 | 5.49749037 | 4.824447397 | 4.713788126 | 5.47077881 | 5.87347892 | 5.50859267 | 5.84037943 | 5.632916569 | 5.56603558 |
| ENSMUSG00000( | Ggta1 | -7.11E-01 | -1.05E+00 | -3.70E-01 | 4.718244 | -4.39E+00 | 3.79E-04 | 3.36E-02 | -0.07465 | -1.63657 | 4.346670705 | 4.79202786 | 4.273178511 | 4.126129503 | 4.62319011 | 4.98076266 | 5.177925045 | 5.112645 | 4.548046737 | 5.20186412 |
| ENSMUSG00000( | Tyrobp | 1.13E+00 | 5.87E-01 | 1.67E+00 | 2.620175 | 4.38E+00 | 3.91E-04 | 3.38E-02 | 0.189352 | 2.189562 | 3.040009367 | 2.75706525 | 2.903167418 | 3.562862073 | 3.22730976 | 2.18928786 | 1.652818944 | 1.86438882 | 2.644821612 | 2.36001553 |
| ENSMUSG00000( | Acsbg1 | -3.99E+00 | -5.91E+00 | -2.07E+00 | -1.24753 | -4.38E+00 | 3.91E-04 | 3.38E-02 | -0.57387 | -15.9017 | -4.26681184 | -3.8248885 | 0.382185543 | -3.35122402 | -5.99427736 | 1.25941395 | 1.353805241 | 0.24145847 | 1.649805019 | 0.07520061 |
| ENSMUSG00000( | Tubb4b | -5.43E-01 | -8.05E-01 | -2.82E-01 | 8.327725 | -4.38E+00 | 3.86E-04 | 3.38E-02 | -0.46667 | -1.45748 | 8.374308154 | 7.9463616 | 7.677443097 | 8.066782455 | 7.94732051 | 8.47380824 | 8.650385009 | 8.56624147 | 9.129837207 | 8.44476126 |
| ENSMUSG00000( | Insl6 | 9.08E-01 | 4.72E-01 | 1.34E+00 | 4.378471 | 4.38E+00 | 3.86E-04 | 3.38E-02 | -0.07677 | 1.876318 | 4.506876864 | 4.34403428 | 5.297614764 | 5.062403905 | 5.08187017 | 4.39859183 | 3.71710377 | 3.68329603 | 3.554704317 | 4.13821041 |
| ENSMUSG00000( | Tnfrsf25 | -3.64E+00 | -5.39E+00 | -1.88E+00 | 0.266164 | -4.37E+00 | 3.94E-04 | 3.39E-02 | -0.29947 | -12.4243 | -5.85177434 | -1.01753358 | -0.192929172 | -0.71895581 | 0.69222317 | 2.06224867 | 2.211484701 | 1.66369147 | 2.231447161 | 1.58173647 |
| ENSMUSG00000( | Gapt | 1.64E+00 | 8.41E-01 | 2.43E+00 | 1.006709 | 4.34E+00 | 4.25E-04 | 3.63E-02 | 0.151164 | 3.105884 | 1.799277355 | 1.46051372 | 1.626989212 | 2.511723224 | 1.58315147 | 0.35120643 | 0.396873962 | -0.31493488 | -0.10383932 | 0.75613217 |
| ENSMUSG00000( | Fcgrt | -6.70E-01 | -9.98E-01 | -3.43E-01 | 6.105082 | -4.31E+00 | 4.54E-04 | 3.83E-02 | -0.46855 | -1.59155 | 5.320027952 | 6.08410449 | 5.650254821 | 5.617442769 | 6.25276006 | 6.55985837 | 6.122871678 | 6.51820325 | 6.622497673 | 6.30279746 |
| ENSMUSG00000( | Usp44 | 1.00E+00 | 5.12E-01 | 1.50E+00 | 2.716734 | 4.30E+00 | 4.68E-04 | 3.83E-02 | 0.004999 | 2.005513 | 3.534088064 | 3.26892517 | 3.308704945 | 3.016845853 | 2.61674744 | 2.47879448 | 2.211484701 | 2.02944806 | 2.703515606 | 1.9987875 |
| ENSMUSG00000( | Itgb3 | -9.46E-01 | -1.41E+00 | -4.83E-01 | 4.530964 | -4.30E+00 | 4.65E-04 | 3.83E-02 | -0.23234 | -1.92678 | 4.278796226 | 4.56828741 | 3.852356876 | 3.979692854 | 3.27484932 | 4.48893451 | 5.207256577 | 5.10341383 | 5.370007992 | 5.18604638 |
| ENSMUSG00000( | Plk2 | -1.27E+00 | -1.89E+00 | -6.46E-01 | 4.750711 | -4.30E+00 | 4.60E-04 | 3.83E-02 | -0.21135 | -2.405 | 2.990576007 | 4.69788917 | 3.752693818 | 3.78422876 | 6.03466469 | 5.53370282 | 5.26186118 | 5.34523583 | 5.281270623 | 4.82499025 |
| ENSMUSG00000( | Slc6a13 | 7.71E-01 | 3.93E-01 | 1.15E+00 | 4.194307 | 4.30E+00 | 4.65E-04 | 3.83E-02 | -0.22186 | 1.706973 | 4.289055434 | 4.61223934 | 4.33829313 | 4.933253094 | 4.25720505 | 3.87734386 | 3.536304935 | 3.8119214 | 4.390248784 | 3.89720231 |
| ENSMUSG00000( | Med12l | -6.40E-01 | -9.53E-01 | -3.26E-01 | 4.766526 | -4.30E+00 | 4.66E-04 | 3.83E-02 | -0.28944 | -1.55805 | 4.207732368 | 4.49246984 | 4.81942058 | 4.425880026 | 4.32976814 | 5.29076793 | 4.99115553 | 5.05503788 | 4.933031989 | 5.11999716 |
| ENSMUSG00000( | Efcc1 | -8.99E-01 | -1.34E+00 | -4.59E-01 | 3.655645 | -4.30E+00 | 4.63E-04 | 3.83E-02 | -0.07099 | -1.86486 | 2.530979365 | 3.48777667 | 3.695908489 | 3.103544095 | 3.27016524 | 4.1886244 | 4.063710739 | 3.9976278 | 4.15478478 | 4.06332982 |
| ENSMUSG00000( | Gsap | -6.58E-01 | -9.82E-01 | -3.35E-01 | 5.165719 | -4.29E+00 | 4.72E-04 | 3.85E-02 | -0.36682 | -1.5783 | 4.598405911 | 5.11233817 | 4.630113056 | 4.770827424 | 5.15864122 | 5.38598655 | 5.495773012 | 5.68550705 | 5.617736823 | 5.20186412 |
| ENSMUSG00000( | Spaca6 | -1.05E+00 | -1.56E+00 | -5.32E-01 | 3.39895 | -4.29E+00 | 4.79E-04 | 3.87E-02 | -0.08088 | -2.06425 | 2.419688691 | 3.0504001 | 3.239484896 | 3.384839604 | 2.45280587 | 4.15148583 | 4.267093607 | 3.99705824 | 3.254530775 | 3.77211296 |
| ENSMUSG00000( | Cbx7 | -8.81E-01 | -1.31E+00 | -4.48E-01 | 4.530982 | -4.28E+00 | 4.84E-04 | 3.88E-02 | -0.2527 | -1.84191 | 3.485847565 | 4.43908485 | 4.351551167 | 3.586591145 | 4.97510916 | 4.97114999 | 4.844686498 | 4.9892039 | 4.980897395 | 4.68569821 |
| ENSMUSG00000( | Ahnak | -1.60E+00 | -2.39E+00 | -8.12E-01 | 3.011248 | -4.28E+00 | 4.83E-04 | 3.88E-02 | 0.002538 | -3.029 | 1.424350069 | 2.50064183 | 2.818069827 | 2.061557501 | 2.67460763 | 3.7063463 | 4.361435682 | 3.82802307 | 3.729639193 | 3.00781377 |
| ENSMUSG00000( | Ifi44 | -6.40E-01 | -9.56E-01 | -3.25E-01 | 5.800633 | -4.27E+00 | 4.90E-04 | 3.90E-02 | -0.50948 | -1.55887 | 5.136200188 | 5.96334042 | 5.569029046 | 5.297672783 | 5.5482709 | 6.29484521 | 5.897750982 | 6.15059151 | 5.945122707 | 6.20350187 |
| ENSMUSG00000( | Apol7e | 9.66E-01 | 4.90E-01 | 1.44E+00 | 4.415593 | 4.27E+00 | 4.93E-04 | 3.90E-02 | -0.33983 | 1.953007 | 4.834848744 | 5.24598379 | 4.764606448 | 5.400119182 | 3.69861707 | 4.41286428 | 3.616584602 | 4.19244937 | 4.036292519 | 3.95356786 |
| ENSMUSG00000( | Eif2s2 | 4.39E-01 | 2.22E-01 | 6.57E-01 | 6.968683 | 4.26E+00 | 5.07E-04 | 3.99E-02 | -0.69281 | 1.355946 | 7.295818563 | 6.97556817 | 7.408197294 | 7.234174225 | 7.00053215 | 6.86683283 | 6.770800525 | 6.67629572 | 6.627577602 | 6.83103349 |
| ENSMUSG00000( | Meis3 | -2.17E+00 | -3.25E+00 | -1.09E+00 | 1.866059 | -4.25E+00 | 5.18E-04 | 4.03E-02 | -0.02549 | -4.50405 | 0.5576166 | 1.12930781 | -1.28407706 | 1.221665644 | 2.08787168 | 2.71212865 | 3.351617681 | 2.3657866 | 3.847106459 | 2.67156788 |
| ENSMUSG00000( | Pbx1 | -6.14E-01 | -9.18E-01 | -3.10E-01 | 6.720425 | -4.25E+00 | 5.18E-04 | 4.03E-02 | -0.67985 | -1.53043 | 6.546167635 | 6.24308358 | 7.013138285 | 6.210636684 | 6.13982804 | 6.84240061 | 7.13707139 | 6.98148654 | 6.807796532 | 7.2826411 |
| ENSMUSG00000( | Slc17a8 | 1.74E+00 | 8.76E-01 | 2.60E+00 | 2.404249 | 4.25E+00 | 5.22E-04 | 4.04E-02 | -0.05553 | 3.3355 | 2.852129237 | 3.44750801 | 3.741182892 | 2.945233383 | 3.03638978 | 2.60017372 | 0.009850839 | 1.92552911 | 1.764516678 | 1.71997251 |
| ENSMUSG00000( | Etfb | 5.34E-01 | 2.68E-01 | 7.99E-01 | 8.193884 | 4.23E+00 | 5.38E-04 | 4.10E-02 | -0.79896 | 1.447676 | 8.624831388 | 8.20612341 | 8.111131069 | 8.691349184 | 8.2618576 | 7.94243415 | 7.882410655 | 7.79710082 | 8.511625229 | 7.90997763 |
| ENSMUSG00000( | Tspan3 | -4.98E-01 | -7.45E-01 | -2.50E-01 | 7.438464 | -4.23E+00 | 5.38E-04 | 4.10E-02 | -0.76582 | -1.41193 | 7.270891912 | 7.33157385 | 7.148192488 | 6.92466767 | 7.36835142 | 7.79358191 | 7.672998368 | 7.70732413 | 7.798991488 | 7.36806999 |
| ENSMUSG00000( | Rbp1 | -6.39E-01 | -9.57E-01 | -3.21E-01 | 6.199831 | -4.23E+00 | 5.35E-04 | 4.10E-02 | -0.64956 | -1.55734 | 5.772564209 | 5.51896343 | 6.146193666 | 5.891473744 | 5.75684823 | 6.64947304 | 6.493046503 | 6.26914334 | 7.219974792 | 6.2806254 |
| ENSMUSG00000( | B4galnt1 | 5.99E-01 | 3.00E-01 | 8.97E-01 | 5.770444 | 4.22E+00 | 5.47E-04 | 4.14E-02 | -0.65167 | 1.514433 | 6.489800081 | 5.90326331 | 5.619684305 | 6.199499678 | 6.01379405 | 5.58977345 | 5.320560089 | 5.6044052 | 5.493081962 | 5.47057901 |
| ENSMUSG00000( | Cep85 | -5.94E-01 | -8.92E-01 | -2.97E-01 | 6.16999 | -4.21E+00 | 5.64E-04 | 4.25E-02 | -0.70897 | -1.50977 | 6.165047877 | 6.05705137 | 5.712910916 | 5.708498976 | 5.99501069 | 6.22796892 | 6.677053016 | 6.6040546 | 6.036149799 | 6.51615808 |
| ENSMUSG00000( | Uqcc2 | 5.35E-01 | 2.67E-01 | 8.02E-01 | 5.967472 | 4.21E+00 | 5.68E-04 | 4.26E-02 | -0.70948 | 1.448548 | 6.248559482 | 5.76569855 | 6.252043431 | 6.331419496 | 6.46027899 | 5.60794828 | 5.713275298 | 5.52290945 | 6.02496432 | 5.74762595 |
| ENSMUSG00000( | Hif3a | -1.31E+00 | -1.97E+00 | -6.54E-01 | 2.560372 | -4.20E+00 | 5.82E-04 | 4.33E-02 | -0.13123 | -2.4859 | 2.108227595 | 2.31670778 | 1.449748827 | 1.095032206 | 2.71652908 | 2.9911046 | 3.279804659 | 3.07603486 | 3.70111711 | 2.86941016 |
| ENSMUSG00000( | Fhdc1 | -1.30E+00 | -1.95E+00 | -6.45E-01 | 2.579397 | -4.19E+00 | 5.86E-04 | 4.34E-02 | -0.14766 | -2.45558 | 0.806437146 | 2.75102886 | 1.940553833 | 2.061557501 | 2.10901045 | 3.10096108 | 3.24722629 | 3.06520783 | 3.267557455 | 3.44443442 |
| ENSMUSG00000( | Mrps10 | 4.27E-01 | 2.12E-01 | 6.42E-01 | 6.155192 | 4.19E+00 | 5.92E-04 | 4.36E-02 | -0.77649 | 1.344344 | 6.347499249 | 6.37354672 | 6.450838687 | 6.353585706 | 6.48798608 | 5.93637478 | 6.066547741 | 5.95272576 | 5.694247378 | 5.88857056 |
| ENSMUSG00000( | Hspe1 | 5.66E-01 | 2.81E-01 | 8.51E-01 | 6.972031 | 4.19E+00 | 5.94E-04 | 4.36E-02 | -0.84997 | 1.480403 | 7.328927396 | 6.71611516 | 7.46722539 | 7.43461747 | 6.93548036 | 6.74315207 | 6.57364258 | 6.59702922 | 7.215477823 | 6.70864566 |
| ENSMUSG00000( | Nrk | -1.97E+00 | -2.96E+00 | -9.76E-01 | 2.685865 | -4.17E+00 | 6.11E-04 | 4.46E-02 | -0.17501 | -3.91844 | 2.342982518 | 2.00800151 | 2.473131754 | 0.108207595 | 2.79688553 | 3.31366671 | 3.907835882 | 4.15857192 | 2.604322249 | 3.1450446 |
| ENSMUSG00000( | Mrto4 | 4.96E-01 | 2.45E-01 | 7.46E-01 | 5.849849 | 4.16E+00 | 6.28E-04 | 4.51E-02 | -0.79441 | 1.409866 | 6.302702041 | 5.967212 | 5.982904977 | 5.980857293 | 6.28844622 | 5.6110464 | 5.655029263 | 5.6268803 | 5.745380287 | 5.33803224 |
| ENSMUSG00000( | Rpl36al | 5.77E-01 | 2.85E-01 | 8.69E-01 | 7.241638 | 4.16E+00 | 6.29E-04 | 4.51E-02 | -0.91932 | 1.491984 | 7.350566855 | 7.20413639 | 7.831195077 | 7.573573637 | 7.29552253 | 7.07429365 | 6.819205915 | 6.72770516 | 7.68645083 | 6.85373463 |
| ENSMUSG00000( | Lratd2 | -7.12E-01 | -1.07E+00 | -3.51E-01 | 4.568268 | -4.16E+00 | 6.28E-04 | 4.51E-02 | -0.55316 | -1.6376 | 4.195022273 | 4.41405118 | 4.611752396 | 4.11122099 | 4.01244143 | 5.22279356 | 5.027937148 | 4.95414228 | 4.258418518 | 4.87489648 |
| ENSMUSG00000: | Gm36963 | -3.35E+00 | -5.04E+00 | -1.65E+00 | -0.54436 | -4.16E+00 | 6.31E-04 | 4.51E-02 | -0.86442 | -10.1646 | -1.94488374 | -3.33946167 | -1.28407706 | -4.08818962 | 1.45693375 | 0.76129471 | 1.167392116 | 1.00132446 | 0.619643049 | 0.20644515 |
| ENSMUSG00000( | Dnah1 | -1.05E+00 | -1.59E+00 | -5.19E-01 | 2.975449 | -4.15E+00 | 6.36E-04 | 4.51E-02 | -0.28397 | -2.07576 | 2.456564694 | 2.74496711 | 1.967148043 | 2.765639734 | 2.11946481 | 3.6592337 | 3.213895219 | 3.74904171 | 3.367741385 | 3.71078919 |
| ENSMUSG00000( | H2-Ab1 | 9.07E-01 | 4.47E-01 | 1.37E+00 | 6.834002 | 4.15E+00 | 6.38E-04 | 4.51E-02 | -0.90886 | 1.874907 | 7.037919743 | 7.4835643 | 7.031761633 | 7.354235575 | 7.08903575 | 7.08675086 | 6.189912992 | 6.02770631 | 7.202699323 | 5.83643522 |
| ENSMUSG00000( | Sfxn1 | 4.22E-01 | 2.07E-01 | 6.37E-01 | 7.452479 | 4.13E+00 | 6.66E-04 | 4.66E-02 | -0.98798 | 1.339722 | 7.877315282 | 7.45808406 | 7.569978232 | 7.657344659 | 7.54621397 | 7.20935778 | 7.155685033 | 7.23347152 | 7.590943718 | 7.22639841 |
| ENSMUSG00000( | Zmat3 | 1.33E+00 | 6.52E-01 | 2.01E+00 | 2.386162 | 4.14E+00 | 6.63E-04 | 4.66E-02 | -0.28549 | 2.513313 | 2.79568409 | 3.67495739 | 3.047229367 | 2.937872678 | 2.36766642 | 0.87562738 | 2.340629852 | 1.51248566 | 2.664652783 | 1.64481515 |
| ENSMUSG00000( | Acsl3 | -8.83E-01 | -1.33E+00 | -4.33E-01 | 5.143414 | -4.13E+00 | 6.68E-04 | 4.66E-02 | -0.70078 | -1.84479 | 4.823300584 | 5.14917039 | 5.135183349 | 4.082603411 | 4.78626241 | 5.70366894 | 5.869919743 | 5.64233335 | 5.072464899 | 5.16923556 |
| ENSMUSG00000( | Acox2 | -4.25E+00 | -6.42E+00 | -2.08E+00 | -1.51235 | -4.13E+00 | 6.73E-04 | 4.67E-02 | -0.90668 | -19.0396 | -4.26681184 | -6.1468166 | -5.807639016 | 0.736238817 | -4.40931486 | 1.64519764 | 1.085515109 | 0.38767921 | 0.659720488 | 0.99273845 |
| ENSMUSG00000( | Lamb3 | 1.28E+00 | 6.25E-01 | 1.93E+00 | 2.060677 | 4.12E+00 | 6.84E-04 | 4.72E-02 | -0.29299 | 2.425317 | 1.962006855 | 2.85599842 | 3.015728224 | 3.278132596 | 2.33215213 | 1.38055039 | 1.666963126 | 1.73373897 | 1.388759166 | 0.99273845 |
| ENSMUSG00000( | Atp5e | 7.19E-01 | 3.51E-01 | 1.09E+00 | 6.68493 | 4.11E+00 | 6.97E-04 | 4.76E-02 | -0.98379 | 1.646243 | 6.835820053 | 6.2575267 | 7.353335359 | 7.136816928 | 7.3903715 | 6.44938276 | 6.223796253 | 5.98526064 | 7.026641455 | 6.19034751 |
| ENSMUSG00000( | Rps27l | 6.18E-01 | 3.02E-01 | 9.34E-01 | 6.522446 | 4.11E+00 | 6.95E-04 | 4.76E-02 | -0.9707 | 1.534655 | 6.835382608 | 6.22686424 | 7.092276748 | 6.903096011 | 6.95973715 | 6.42494451 | 6.104801452 | 5.89947948 | 6.453245857 | 6.32463391 |
| ENSMUSG00000( | Mamdc2 | -6.24E-01 | -9.43E-01 | -3.04E-01 | 5.431689 | -4.11E+00 | 7.00E-04 | 4.76E-02 | -0.80043 | -1.54087 | 5.014731876 | 5.1924767 | 5.119398153 | 4.79953872 | 5.74339251 | 5.77378966 | 5.642566472 | 5.88030463 | 5.731403238 | 5.41928512 |
| ENSMUSG00000( | Ndufa12 | 4.86E-01 | 2.36E-01 | 7.36E-01 | 6.824863 | 4.10E+00 | 7.20E-04 | 4.82E-02 | -1.02959 | 1.400669 | 6.966879734 | 6.60438851 | 7.220149515 | 7.147318066 | 7.32341789 | 6.64403355 | 6.487094755 | 6.52482715 | 6.859339055 | 6.47117738 |
| ENSMUSG00000( | Epb41 | -4.94E-01 | -7.48E-01 | -2.40E-01 | 6.458885 | -4.10E+00 | 7.17E-04 | 4.82E-02 | -0.98017 | -1.40832 | 6.16538205 | 6.54120636 | 6.320677587 | 5.962819879 | 6.15260948 | 6.54966829 | 6.642781182 | 6.79584267 | 6.626295074 | 6.83157161 |
| ENSMUSG00000( | Tacstd2 | 6.34E-01 | 3.08E-01 | 9.59E-01 | 5.523027 | 4.10E+00 | 7.17E-04 | 4.82E-02 | -0.89048 | 1.551679 | 5.9632086 | 5.97504065 | 5.806610712 | 5.949357336 | 5.46156355 | 5.23886522 | 4.952545783 | 5.37403531 | 4.821939355 | 5.68710366 |
| ENSMUSG00000( | Myl10 | 1.97E+00 | 9.56E-01 | 2.99E+00 | 6.802132 | 4.08E+00 | 7.42E-04 | 4.95E-02 | -1.05409 | 3.929103 | 7.425948945 | 8.59628219 | 7.85647489 | 8.183954133 | 5.69790222 | 6.80906513 | 5.024343473 | 4.72774379 | 6.245917644 | 7.45368471 |
| ENSMUSG00000( | Ltv1 | 4.83E-01 | 2.33E-01 | 7.32E-01 | 6.028331 | 4.08E+00 | 7.53E-04 | 4.98E-02 | -1.00964 | 1.397529 | 6.442559223 | 6.30876749 | 6.512879508 | 6.326495618 | 5.83821713 | 5.84481516 | 5.897001141 | 5.90480309 | 5.323985109 | 5.8837818 |
| ENSMUSG00000( | Rnf208 | -1.83E+00 | -2.77E+00 | -8.84E-01 | 1.935177 | -4.08E+00 | 7.54E-04 | 4.98E-02 | -0.34706 | -3.55326 | 1.067088901 | 0.36097804 | -0.253050164 | 1.842547719 | 1.75391549 | 2.16700642 | 3.012622589 | 2.60135441 | 3.544037392 | 3.25526762 |

1. Greenberg ZJ, Paracatu LC, Monlish DA, et al. The tetraspanin CD53 protects stressed hematopoietic stem cells via promotion of DREAM complex-mediated quiescence. *Blood*. 2023;141(10):1180-1193.
2. Lam J, Katti P, Biete M, et al. A Universal Approach to Analyzing Transmission Electron Microscopy with ImageJ. *Cells*. 2021;10(9).
